# Phylogenomics of Ichneumonoidea (Hymenoptera) and implications for evolution of mode of parasitism and viral endogenization

**DOI:** 10.1101/2020.06.17.157719

**Authors:** Barbara J. Sharanowski, Ryan D. Ridenbaugh, Patrick K. Piekarski, Gavin R. Broad, Gaelen R. Burke, Andrew R. Deans, Alan R. Lemmon, Emily C. Moriarty Lemmon, Gloria J. Diehl, James B. Whitfield, Heather M. Hines

**Author notes:** **Corresponding Author:** Barbara Sharanowski.

## Abstract

Ichneumonoidea is one of the most diverse lineages of animals on the planet with more than 48,000 described species and many more undescribed. Parasitoid wasps of this superfamily are beneficial insects that attack and kill other arthropods and are important for understanding diversification and the evolution of life history strategies related to parasitoidism. Further, some lineages of parasitoids within Ichneumonoidea have acquired endogenous virus elements (EVEs) that are permanently a part of the wasp’s genome and benefit the wasp through host immune disruption and behavioral control. Unfortunately, understanding the evolution of viral acquisition, parasitism strategies, diversification, and host immune disruption mechanisms, is deeply limited by the lack of a robust phylogenetic framework for Ichneumonoidea. Here we design probes targeting 541 genes across 91 taxa to test phylogenetic relationships, the evolution of parasitoid strategies, and the utility of probes to capture polydnavirus genes across a diverse array of taxa. Phylogenetic relationships among Ichneumonoidea were largely well resolved with most higher-level relationships maximally supported. We noted codon use biases between the outgroups, Braconidae, and Ichneumonidae and within Pimplinae, which were largely solved through analyses of amino acids rather than nucleotide data. These biases may impact phylogenetic reconstruction and caution for outgroup selection is recommended. Ancestral state reconstructions were variable for Braconidae across analyses, but consistent for reconstruction of idiobiosis/koinobiosis in Ichneumonidae. The data suggest many transitions between parasitoid life history traits across the whole superfamily. The two subfamilies within Ichneumonidae that have polydnaviruses are supported as distantly related, providing strong evidence for two independent acquisitions of ichnoviruses. Polydnavirus capture using our designed probes was only partially successful and suggests that more targeted approaches would be needed for this strategy to be effective for surveying taxa for these viral genes. In total, these data provide a robust framework for the evolution of Ichneumonoidea.

## 1. Introduction

Ichneumonoidea is one of 15 superfamilies of parasitoid Hymenoptera, wasps whose larvae feed on other arthropods. As a result of an exceptional rate of species diversification, this lineage comprises ~48,000 valid species, 33% of all Hymenoptera (145,000 spp.; (Huber, 2009)) and 3% of all known life (Chapman, 2009; Yu et al., 2016). The true diversity may be many times higher when undescribed species are considered. For example, estimates of known and unknown diversity in a single polydnavirus-carrying subfamily (Microgastrinae) within the group might exceed the total described species for the superfamily (Rodriguez et al., 2013). Ichneumonoidea has been challenging for higher level systematics because of its rapid radiation and relatively high levels of morphological convergence (Quicke, 2015; Sharkey, 2007). Further, the sheer size of the lineage and the relative dearth of taxonomic experts in this group has presented many challenges for large phylogenetic studies.

The diversity of Ichneumonoidea may be due to the varied methods the wasps use to attack and kill their hosts as the evolution of new mechanisms to exploit hosts may have opened up new niches, enabling adaptive radiations (Price, 1980). Wasp species may be external (ectoparasitic) or internal (endoparasitic) parasitoids, although the latter often leave the host to pupate (Austin and Dowton, 2000). Some ichneumonoids are idiobionts, arresting the development of their host upon oviposition; whereas others are koinobionts, allowing the host to continue development, often allowing the host to molt to other life stages (Askew and Shaw, 1986). Further, ichneumonoids attack their arthropod hosts in all life stages, from eggs to adults, and hosts in different habitats, from exposed to concealed (Quicke, 2015).

Understanding core phylogenetic relationships in Ichneumonoidea is pertinent to research across multiple fields. Ichneumonoid parasitoid wasps are beneficial insects, providing natural management of pests in crops and forests, and are estimated to provide >$17 billion in these ecosystem services (Pimentel et al., 1997). Many ichneumonoids have been released in biological control programs against major invasive pests, such as the Emerald ash borer (Bauer et al., 2008; Kula et al., 2010) and the Mediterranean fruit fly (Vargas et al., 2001). Several ichneumonoid wasps are important models for understanding parasitoid physiology (Ballesteros et al., 2017; Webb et al., 2006), interactions with host immune response (Lovallo et al., 2002; Schmidt et al., 2001; Strand, 2012), development of novel transgenic control methods for insect pests (Beckage and Gelman, 2004; Webb et al., 2017), and the process and evolution of viral endogenization to produce beneficial viruses (Bézier et al., 2009; Burke et al., 2018a; Pichon et al., 2015; Strand and Burke, 2012). Given their immense diversity, Ichneumonoidea also present an exceptional opportunity to understand the factors that promote diversification, a critically important question as we face increasing species extinctions with continued habitat destruction and global climate change.

### 1.1 Phylogenetic history of Ichneumonoidea

Ichneumonoidea is robustly monophyletic (Sharanowski et al., 2011; Sharkey et al., 2010) and currently comprises the monophyletic families: Ichneumonidae, Braconidae and Trachypetidae (Quicke et al., 2020b; Sharkey, 2007). Previously in Braconidae, Trachypetidae was recently elevated to family status (Quicke et al., 2020b), however the data supporting this taxonomic status is not robust, and as we do not sample this lineage here, we do not discuss it further. The classification of the other two families has also been in flux, with Braconidae having anywhere from 17 to 50 subfamilies and Ichneumonidae with 24 to 44 subfamilies (Belshaw and Quicke, 2002; Bennett et al., 2019; Quicke et al., 2020a). Braconidae, represented by over 21,000 valid species (Yu et al., 2016), 41 currently recognized subfamilies (Sharanowski et al., 2011), and approximately 109 tribes, are highly valued for their use in biocontrol of pestiferous hosts (Austin and Dowton, 2000; Peixoto et al., 2018; Zhang et al., 2017). A molecular phylogeny of Braconidae using 7 gene regions was completed by Sharanowski et al. (2011). Hypotheses of deep level relationships were well supported, allowing for a revised classification, but relationships within one of the three major lineages (cyclostomes) remain largely unresolved (Sharanowski et al., 2011; Zaldivar-Riverón et al., 2006), and will require greater taxon representation to allow more comprehensive reclassification.

Although Ichneumonidae is currently larger in terms of number of described species than Braconidae, the family is much less understood phylogenetically. Ichneumonidae is comprised of over 25,000 valid species, 38-43 subfamilies, and approximately 57 tribes (Bennett et al., 2019; Broad et al., 2018; Quicke et al., 2020a; Quicke, 2015; Quicke et al., 2009; Yu et al., 2012). Until recently, the three molecular studies that examined higher-level relationships within Ichneumonidae (Belshaw et al., 1998; Quicke et al., 2000; Quicke et al., 2009) were all based on one gene (28S rDNA), severely limiting phylogenetic resolution. Phylogenetic and revisionary studies have been attempted for some lineages (e.g. Bennett, 2001; Gauld et al., 2002; Gauld, 1985; Wahl and Gauld, 1998), with only a few based on molecular data (Klopfstein et al., 2010; e.g. Laurenne et al., 2006; Matsumoto, 2016; Rousse et al., 2016; Santos, 2017). There are three informal lineages in Ichneumonidae that have been recognized for many years, namely Ichneumoniformes, Ophioniformes and Pimpliformes (Wahl, 1986, 1991; Wahl, 1993a; Wahl and Gauld, 1998). These definitions were largely kept but redefined in 2009 with Quicke et al.’s (2009) combined 28S and morphological analyses based on dense taxonomic sampling (1001 wasps). Quicke et al. (2000) also added three additional informal groupings for single subfamily lineages (Xoridiformes, Labeniformes, Orthopelmatiformes), the latter two with uncertain placement, even when the taxonomic sampling was robust (Quicke et al., 2009). This classification, based on a single gene, was largely followed in Quicke’s (2015) comprehensive and seminal book on Ichneumonoidea.

Several recent studies have been published with more comprehensive taxon, genetic, and morphological sampling. Santos (2017) examined relationships among Cryptinae and higher-level relationships within Ichneumoniformes using extensive taxon sampling, morphological characters, and seven genes and revised the taxonomy for some lineages. Klopfstein et al. (2019) analyzed relationships among the Pimpliformes using anchored hybrid enrichment with 93 genes and 723 genes from a reduced taxon transcriptome dataset. They showed the evolutionary history of Pimpliformes was confounded by rapid radiations across several lineages. Finally, Bennett et al. (2019) examined relationships among the whole family using extensive morphological characters and three loci, and currently represents the most detailed analysis and summary of ichneumonid relationships to date. Here we follow the revised higher-level groupings for Ichneumonidae proposed by Bennett et al. (2019).

Given the immense diversity on both Braconidae and Ichneumonidae, continued research on their phylogenetic relationships, evolution, and taxonomy are warranted given their importance in pest control and interesting evolutionary history with endogenous viruses. Further, members of both families have been underutilized in biological control due to numerous cryptic species (Derocles et al., 2016; Peixoto et al., 2018) and limited taxonomic resources facilitating accurate identification of many groups, particularly for species (Broad et al., 2018; Quicke, 2015; Veijalainen et al., 2012; Wahl, 1993b).

### 1.2 Evolution of viral symbiosis in Ichneumonoidea

Some ichneumonoids overcome host defenses through the use of viruses produced by genes that are integrated within the wasp genome and modify host physiology and immunity (Kroemer and Webb, 2004; Lavine and Beckage, 1995; Strand and Burke, 2012). The acquisition of these endogenous virus elements (EVEs) in Ichneumonoidea promotes reproductive success and may be a factor promoting diversity (Burke et al., 2018a; Burke and Strand, 2014; Federici and Bigot, 2003; Gauthier et al., 2017; Whitfield, 1990). These viruses in the family Polydnaviridae (PDV, genera *Bracovirus*, *Ichnovirus*) are only known from a few clades across Ichneumonoidea. Within Ichneumonidae, viruses are known from two subfamilies (Banchinae and Campopleginae) that are both members of the Ophioniformes (Quicke, 2015; Quicke et al., 2009). In both lineages, the viruses are ichnoviruses (PDVs) that have some homologous genes, but whether this represents one or two independent acquisitions remains unclear (Béliveau et al., 2015; Lapointe et al., 2007; Volkoff et al., 2012; Volkoff et al., 2010), largely due to a lack of knowledge on Ichneumonid relationships and viral presence across the family. There is one known case of an ichneumonid, *Venturia canescens* (Campopleginae), which does not produce an ichnovirus but instead has an alphanudivirus-like EVE that no longer packages DNA but produces virus-like particles (VLPs) (Pichon et al. 2015). Thus, for Ichneumonidae, there have been at least 2 or 3 independent acquisitions of EVEs.

Within Braconidae, EVEs are best known within the diverse microgastroid complex, in which all known viruses are bracoviruses (Polydnaviridae) that were likely derived from a common ancient (~100 mya) betanudivirus (Nudiviridae) (Bézier et al., 2009; Bézier et al., 2013; Murphy et al., 2008; Whitfield, 1997, 2002; Whitfield and O’Connor, 2012). The latest discovery of an independent and recent acquisition of endogenous virus related to alphanudiviruses (FaENV - Nudiviridae) in a different lineage of Braconidae (some species of *Fopius* (Opiinae)) suggests that viral endogenization may be more common than previously thought (Burke et al., 2018a). Unlike bracoviruses and ichnoviruses, FaENV in species of *Fopius*, like in *Venturia*, lost the ability to package DNA and instead produce VLPs. Thus, both Ichneumonidae and Braconidae have at least two subfamilies that produce viruses (PDVs and VLPs), but virion morphology, virus replication machinery, genome rearrangement and gene homology suggest these endogenization events likely represent four or five independent acquisitions (Béliveau et al., 2015; Burke et al., 2018a; Herniou et al., 2013; Thézé et al., 2011).

### 1.3 Importance of a stable phylogenetic framework for Ichneumonoidea

Unfortunately, understanding of viral acquisition and evolution, as well as parasitoid evolution, host use, development, diversification, and host immune disruption, is severely limited by the lack of a robust phylogenetic framework (Desjardins et al., 2008; Gauld, 1988; Lapointe et al., 2007; Pennacchio and Strand, 2006; Whitfield and O’Connor, 2012). Outside of low genetic sampling of previous studies, the difficulty of resolving clades in Ichneumonoidea is likely due in part to several rapid radiations (Banks and Whitfield, 2006; Klopfstein et al., 2019; Whitfield and Kjer, 2008). Further, previous phylogenetic analyses based on morphology (Belshaw et al., 2003; Dowton et al., 2002; Pitz et al., 2007; Quicke et al., 2000; Sharkey and Wahl, 1992; van Achterberg and Quicke, 1992; Wahl and Gauld, 1998; Wharton et al., 1992) have discovered high levels of convergence, thereby obfuscating true relationships (Gauld and Mound, 1982; Quicke, 2015; Whitfield, 2002; Wild et al., 2013). Thus, deep genetic sampling will likely be necessary to understand relationships in Ichneumonoidea. Several recent studies of other insect groups that likely had rapid radiations have shown the power of a phylogenomics approach using hundreds of genes (Branstetter et al., 2017; Peters et al., 2017; Piekarski et al., 2018), even if taxonomic sampling was relatively low (Johnson et al., 2013; Misof et al., 2014; Savard et al., 2006; Simon et al., 2009).

In this study we resolve several deep phylogenetic relationships for Ichneumonoidea using anchored hybrid enrichment (AHE) (Lemmon et al., 2012) and focusing more on the poorly resolved ichneumonids. We sequenced whole genomes of 11 ichneumonoid wasps and utilize these genomes along with previously sequenced Hymenopteran genomes to design a novel probe set targeted for Ichneumonoidea, but capable of capturing genes across multiple lineages of Hymenoptera. We sequenced 479 genes for 80 ingroup exemplars, one of the largest molecular datasets for this immensely diverse superfamily to date. These data provide a robust phylogenetic framework for Ichneumonoidea and establish molecular tools for future sampling in the lineage. We reconstruct ancestral states for two traits related to parasitism: idiobiosis versus koinobiosis, and ecto- versus endoparasitism. Further we examine the utility of probes to capture polydnaviruses across a broad array of taxa. Our data help resolve previous phylogenetic uncertainties and improve the ability to address interesting biological questions in this rapid radiation.

## 2. METHODS

### 2.1 Genomic Sequencing and Probe Design

The goals for probe design for AHE (Lemmon et al., 2012) were to specifically target ichneumonoids, but also to have broader applicability across all of Hymenoptera. Thus, we utilized published Hymenoptera sequences and sequenced several new genomes to design the probe set. The probe design was generated by optimizing across published genomes from the bees *Apis mellifera* (Honeybee Genome Sequencing Consortium, 2006) and *Bombus impatiens* (Sadd et al., 2015), the parasitoid wasp *Nasonia vitripennis* (Werren et al., 2010), seven ant species (Gadau et al., 2012), and two braconid genomes from i5k (http://arthropodgenomes.org/wiki/i5K): (Microgastrinae: *Microplitis demolitor* (Burke et al., 2018b) and Opiinae: *Diachasma alleoleum* (Tvedte et al., 2019)). To these we added whole genome sequences from 11 additional ichneumonoid wasps, including four braconids and seven ichneumonids that represent diverse lineages across the ichneumonoid phylogeny (Table 1). For whole genome sequencing of these 11 individuals, DNA was extracted from whole specimens using a DNeasy Blood and Tissue Kit with RNase treatment. DNA was only used if good quality scores and DNA concentration were obtained from the Nanodrop, and gel electrophoresis showed minimal DNA degradation. Indexed libraries with ~250 bp inserts were prepared for sequencing following Lemmon et al. (2012), and sequenced at 100bp paired end to achieve ~15x genomic coverage on two HiSeq2500 sequencing lanes (79 Gb total output). Reads were submitted to the NCBI Sequence Read Archive (FigShare 10.6084/m9.figshare.12497468).

**Table 1.** Taxon sampling used in this study, including voucher codes (where appropriate), taxonomy, and country of origin. Genome indicates publicly available genomes used in the study. Bolded taxa are genomes sequenced in this study.

| Outgroups |  |  |  |  |  |
| --- | --- | --- | --- | --- | --- |
| Voucher Code | FSU Code | Family | Subfamily | Genus species | Country |
| N/A | I15712 | Pteromalidae | Pteromalinae | *Nasonia vitripennis* | N/A |
| N/A | I15697 | Apidae | Apinae | *Apis mellifera* | N/A |
| N/A | I15699 | Apidae | Apinae | *Bombus impatiens* | N/A |
| N/A | I15707 | Megachilidae | Megachilinae | *Megachile rotunda* | N/A |
| N/A | I15693 | Formicidae | Myrmicinae | *Acromyrmex echinator* | N/A |
| N/A | I15698 | Formicidae | Myrmicinae | *Atta cephalotes* | N/A |
| N/A | I15700 | Formicidae | Formicinae | *Camponotus floridanus* | N/A |
| N/A | I15705 | Formicidae | Ponerinae | *Harpegnathos saltator* | N/A |
| N/A | I15706 | Formicidae | Dolichoderinae | *Linepithema humile* | N/A |
| N/A | I15715 | Formicidae | Myrmicinae | *Pogonomyrmex barbatus* | N/A |
| N/A | I15716 | Formicidae | Myrmicinae | *Solenopsis invicta* | N/A |
| Braconidae |  |  |  |  |  |
| Voucher Code | FSU Code | Higher-Group | Subfamily | Genus species | Country |
| PSUCFEM10030041 | I2347 | Aphidioid | Aphidiinae | *Pauesia* sp. | Canada: Manitoba |
| PSUCFEM10030046 | I2352 | Aphidioid | Aphidiinae | *Praon* sp. | Canada: Manitoba |
| PSUCFEM10030015 | I2325 | Cyclostomes s.s. | Rhyssalinae | *Rhyssalus* sp. | USA: California |
| **PSUCFEM10030003** | **I15695** | **Cyclostomes s.s.** | **Rogadinae** | ***Aleiodes* sp.** | **Canada: Manitoba** |
| PSUCFEM10030006 | I2358 | Cyclostomes s.s. | Pambolinae | *Pambolus* sp. | USA: Arizona |
| PSUCFEM10030050 | I2355 | Cyclostomes s.s. | Braconinae | *Bracon* sp. | Canada: Manitoba |
| PSUCFEM10030007 | I2359 | Cyclostomes s.s. | Alysiinae | *Chorebus* sp. | Canada: Manitoba |
| Genome | **I15701** | Cyclostomes s.s. | Opiinae | *Diachasma alloeum* | Genome |
| PSUCFEM10030043 | I2349 | Microgastroid | Cardiochilinae | *Schoenlandella* sp. | Thailand: Loei |
| Genome | **I15711** | Microgastroid | Microgastrinae | *Micropilitus demoliter* | Genome |
| PSUCFEM10030045 | I2351 | Microgastroid | Cheloninae | *Chelonus* sp. | Canada: Manitoba |
| **PSUCFEM10030009** | **I15714** | **Microgastroid** | **Cheloninae** | ***Phanerotoma* sp.** | **Costa Rica: Guanacaste** |
| PSUCFEM10030035 | I2343 | Microgastroid | Ichneutinae | *Ichneutes* sp. | Canada: Yukon |
| PSUFEM10030037 | I2344 | Sigalphoid | Agathidinae | *Bassus* sp. | USA: Arizona |
| **PSUCFEM10030008** | **I15710** | **Euphoroid** | **Euphorinae** | ***Meterous* sp.** | **Costa Rica: Guanacaste** |
| PSUCFEM10030032 | I2342 | Helconoid | Orgilinae | *Orgilus ablusus* * | Canada: Manitoba |
| **PSUCFEM10030004** | **I15704** | **Helconoid** | **Helconinae** | ***Eumacrocentrus americanus*** | **USA: West Virginia** |
| PSUCFEM10030049 | I2354 | Helconoid | Acampsohelconinae | *Urosigalphus* sp. | USA: Kansas |
| PSUCFEM10030039 | I2346 | Helconoid | Brachistinae | *Apoblacus* sp. | Chile: X Region |
| PSUCFEM10030038 | I2345 | Helconoid | Brachistinae | *Blacus* sp. | French Guiana: Patawa |
| PSUCFEM10030048 | I2353 | Helconoid | Brachistinae | *Diospilus* sp. | USA: West Virginia |
| Ichneumonidae |  |  |  |  |  |
| **PSUCFEM10030002** | **I15713** | **Xoridiformes** | **Xoridinae** | ***Odontocolon* sp.** | **Canada: Manitoba** |
| PSUCFEM10030023 | I9984 | Orthopelmatiformes | Orthopelmatinae | *Orthopelma* sp. | Canada: Manitoba |
| PSUCFEM000070305 | I9986 | Unknown | Eucerotinae | *Euceros* sp. | Sweden: Uppland |
| PSUCFEM000070330 | I10002 | Labeniformes | Labeninae | *Labium* sp. | Australia: Tasmania |
| PSUCFEM000070316 | I9990 | Labeniformes | Labeninae | *Certonotus nitidulus* | Australia: Tasmania |
| PSUCFEM000070320 | I9994 | Labeniformes | Labeninae | *Certonotus nitidulus* | Australia: Tasmania |
| **PSUCFEM10030001** | **I15694** | **Ichneumoniformes** | **Adelognathinae** | ***Adelognathus* sp.** | **Canada: Manitoba** |
| PSUCFEM10030018 | I2328 | Ichneumoniformes | Cryptinae | *Mesostenus thoracicus* | Canada: Manitoba |
| PSUCFEM10030019 | I2329 | Ichneumoniformes | Cryptinae | *Echthrus abdominalis* | Canada: Manitoba |
| PSUCFEM000070324 | I9997 | Ichneumoniformes | Ichneumoninae | *Gavrana* sp. | Australia: Queensland |
| PSUCFEM000070340 | I10010 | Ichneumoniformes | Ichneumoninae | Ichneumonini (genus ?) | Australia: Tasmania |
| PSUCFEM000070315 | I9989 | Ichneumoniformes | Ichneumoninae | *Akymichneumon* sp. | Australia: Tasmania |
| PSUCFEM000070327 | I9999 | Ichneumoniformes | Ichneumoninae | *Gavrana maculipes* s.g. | Australia: Tasmania |

| Ichneumonidae |  |  |  |  |  |
| --- | --- | --- | --- | --- | --- |
| Voucher Code | FSU Code | Higher Group | Subfamily | Genus species | Country |
| PSUCFEM000070326 | I9998 | Pimpliformes | Pimplinae | *Echthromorpha intricatoria* | Australia: Tasmania |
| PSUCFEM10030020 | I2330 | Pimpliformes | Pimplinae | *Pimpla pedalis* | Canada: Manitoba |
| PSUCFEM000070323 | I9996 | Pimpliformes | Pimplinae | *Pimpla molesta* | Honduras: FM |
| **PSUCFEM10030005** | **I15702** | **Pimpliformes** | **Pimplinae** | ***Dolichomitus* sp.** | **Canada: Manitoba** |
| PSUCFEM10030052 | I2356 | Pimpliformes | Pimplinae | *Scambus* sp. | Canada: Manitoba |
| PSUCFEM000070333 | I10004 | Pimpliformes | Pimplinae | *Eriostethus pulcherrimus* | Australia: Tasmania |
| PSUCFEM10030023 | I2333 | Pimpliformes | Orthocentrinae | *Orthocentrus* sp. | Canada: Manitoba |
| PSUCFEM10030021 | I2331 | Pimpliformes | Poemeniinae | *Neoxorides borealis* | Canada: Manitoba |
| PSUCFEM10030016 | I2326 | Pimpliformes | Acaenitinae | *Spilopteron occiputale* | Canada: Manitoba |
| PSUCFEM000070310 | I9988 | Pimpliformes | Cylloceriainae | *Cylloceria* sp. | Sweden: Halland |
| PSUCFEM000070308 | I9987 | Pimpliformes | Collyriinae | *Collyria* sp. | Sweden: Öland |
| PSUCFEM10030022 | I2332 | Pimpliformes | Diplazontinae | *Homotropus* sp. | Canada: Manitoba |
| PSUCFEM10030014 | I2324 | Ophioniformes | Tryphoninae | *Exenterus* sp. | Canada: Manitoba |
| PSUCFEM10030025 | I2335 | Ophioniformes | Tryphoninae | *Netelia* (*Netelia*) sp. | Canada: Manitoba |
| PSUCFEM000070300 | I9983 | Ophioniformes | Tryphoninae | *Phytodietus* sp. | Sweden: Uppland |
| PSUCFEM10030031 | I2341 | Ophioniformes | Tersilochinae | *Phradis*? | USA: Florida |
| PSUCFEM000070318 | I9992 | Ophioniformes | Tersilochinae | *Stethantyx* sp. | Honduras: FM |
| PSUCFEM10030028 | I2338 | Ophioniformes | Banchinae | *Glypta* sp. | Canada: Manitoba |
| PSUCFEM000070337 | I10008 | Ophioniformes | Banchinae | *Australoglypta* sp. | Australia: Tasmania |
| PSUCFEM000070329 | I10001 | Ophioniformes | Banchinae | *Meniscomorpha* sp.1 | Honduras: FM |
| **PSUCFEM10030012** | **I15708** | **Ophioniformes** | **Banchinae** | ***Lissonota* sp.** | **Canada: Manitoba** |
| PSUCFEM000070332 | I10003 | Ophioniformes | Banchinae | *Meniscomorpha* sp.2 | Honduras: FM |
| PSUCFEM000070348 | I10013 | Ophioniformes | Banchinae | *Lissonota* sp.* | Australia: Tasmania |
| PSUCFEM000070303 | I9985 | Ophioniformes | Oxytorinae | *Oxytorus* sp. | Sweden: Östergötland |
| PSUCFEM000070328 | I10000 | Ophioniformes | Ctenopelmatinae | *Megacera* sp. | Australia: Tasmania |
| **PSUCFEM10030000** | **I15696** | **Ophioniformes** | **Ctenopelmatinae** | ***Anoncus* sp.** | **Canada: Manitoba** |
| PSUCFEM10030030 | I2340 | Ophioniformes | Ctenopelmatinae | *Mesoleius* sp.* | Canada: Manitoba |
| **PSUCFEM10030011** | **I15709** | **Ophioniformes** | **Mesochorinae** | ***Mesochorus* sp.** | **Canada: Manitoba** |
| PSUCFEM10030024 | I2334 | Ophioniformes | Metopiinae | *Exochus* sp. | Canada: Manitoba |
| PSUCFEM10030053 | I2357 | Ophioniformes | Metopiinae | *Spudaeus* sp. | Canada: Manitoba |
| PSUCFEM000070319 | I9993 | Ophioniformes | Metopiinae | *Metopius* sp. | Australia: Tasmania |
| PSUCFEM10030017 | I2327 | Ophioniformes | Ophioninae | *Ophion* sp. | Canada: Manitoba |
| PSUCFEM000070291 | I9982 | Ophioniformes | Hybrizontinae | *Hybrizon buccatus* | Sweden: Gotland |
| PSUCFEM10030026 | I2336 | Ophioniformes | Cremastinae | *Cremastus* sp. | Canada: Manitoba |
| PSUCFEM000070281 | I9981 | Ophioniformes | Cremastinae | *Pristomerus* sp.* | Australia: Tasmania |
| PSUCFEM10030027 | I2337 | Ophioniformes | Anomaloniinae | *Agrypon* sp. | Canada: Manitoba |
| PSUCFEM000070338 | I10009 | Ophioniformes | Anomaloniinae | *Habronyx* sp.* | Australia: Tasmania |
| **PSUCFEM10030013** | **I15703** | **Ophioniformes** | **Campopleginae** | ***Dusona* sp.2** | **Canada: Manitoba** |
| PSUCFEM10030029 | I2339 | Ophioniformes | Campopleginae | *Campoletis* sp.2 | Canada: Manitoba |
| PSUCFEM000070317 | I9991 | Ophioniformes | Campopleginae | *Campoplex* sp.2 | Australia: Tasmania |
| PSUCFEM000070322 | I9995 | Ophioniformes | Campopleginae | *Hyposoter* sp.* | Australia: Tasmania |
| PSUCFEM000070334 | I10005 | Ophioniformes | Campopleginae | *Casinaria* sp.1 | Australia: Tasmania |
| PSUCFEM000070335 | I10006 | Ophioniformes | Campopleginae | *Campoplex* sp.1* | Australia: Tasmania |
| PSUCFEM000070341 | I10011 | Ophioniformes | Campopleginae | *Dusona* sp.3* | Australia: Tasmania |
| PSUCFEM000070336 | I10007 | Ophioniformes | Campopleginae | *Campoletis* sp.1 | Australia: Tasmania |
| PSUCFEM000070347 | I10012 | Ophioniformes | Campopleginae | *Dusona* sp.1 | Australia: Tasmania |

Raw reads from newly sequenced taxa were combined with assembled genome data to design the Hymenoptera probe set. In general, we followed locus selection procedures outlined in Hamilton et al. (2016). Initial Hymenoptera probe design involved finding orthologs to an optimal set of 941 insect loci established in Coleoptera (Haddad et al., 2018). Using the red flour beetle (*Tribolium castaneum*) sequence as a reference, we scanned the assembled genomes of two divergent hymenopteran species (*Bombus impatiens* and *Nasonia vitripennis)* for each of the Coleoptera anchored hybrid enrichment loci. The best matching region in each of the two Hymenoptera genomes was identified and a 4000 bp region containing each locus was extracted and aligned using MAFFT v7.023b (Katoh and Standley, 2013b) with –genafpair and –maxiterate 1000 flags. We then scanned the 12 assembled and raw reads from 11 unassembled genomes outlined above for targeted anchor regions using *Nasonia vitripennis* sequences. For the 12 assembled genomes, we isolated up to 4000 bp surrounding the region that best matched the reference. For the unassembled genomes, reads were merged and trimmed following Rokyta et al. (2012), and then mapped to *Nasonia vitripennis*. The consensus sequences from the resulting assemblies were extended in flanking regions to produce up to a 4000 bp sequence for each species at each locus following protocols outlines in Hamilton et al. (2016). The resulting loci from *N. vitripennis* and the 23 target species were then aligned using MAFFT. These were visually inspected to confirm adequate alignment in Geneious (Kearse et al., 2012), and then trimmed and masked to isolate well-aligned regions. To ensure sufficient enrichment efficiency, we removed loci that contained fewer than 50% of the taxa, resulting in 528 anchor loci.

We also added a few custom selected genes into the probeset: five recognized venom proteins (calreticulin, phenoloxidases, chitinase, trehalase, aspartylglucosaminidase); two genes (ACC and CAD) used in a previous braconid phylogeny (Sharanowski et al., 2011); and five polydnavirus genes (see below). For each of these, rather than use the genomes, probes were designed from a reference set of previously published gene sequences.

Polydnaviruses were included to enable comparison of taxon incidence, number of origins, age of virus endogenization events and evolution of polydnaviruses among the Ichneumonoidea while also assessing the ability of the anchored enrichment approach to capture more rapidly evolving adaptive genes. Furthermore, this capture could increase the numbers of these genes across taxa to enable an improved framework for examining their evolution in future. The PDV genes included two markers identified from bracoviruses (Bézier et al., 2009) from three species: *HzNVorf128*, a gene of viral origin associated with replication, and *p74 per os infectivity factor*, a viral envelope gene. We also designed probes for two ichnovirus replication or structural proteins that occur across three species of Campopleginae, including EST data derived from Sharanowski et al. (2010): IVSP4 and U1 (Burke et al., 2013; Lorenzi et al., 2019; Volkoff et al., 2012; Volkoff et al., 2010). These genes are part of the replication machinery maintained within the host genome and were selected because they are more likely to be single copy or involve just a few copies. In addition, we included sequences representing the vankyrin gene family across all PDV lineages to test the ability to capture gene family representatives (Kroemer and Webb, 2005). Unlike the other loci, vankyrins are packaged within PDVs and are likely not of viral origin (Huguet et al., 2012).

Repetitive Kmer alignment regions among all loci were masked following Hamilton et al. (2016). To build the final probe set we tiled 120 bp probes at 2X density across the 24 taxon-specific sequences for each locus. The final probe set included 57,066 probes from 541 loci covering 212,392 total base pairs. Loci included in the probe design consisted of a diverse array of genes, with no significant enrichment of any particular class of genes based on Blast2GO (Conesa et al., 2005) analysis.

### 2.2 Taxon Sampling

To provide a deeper phylogenetic framework for systematic revision in Ichneumonoidea, we focused our taxon sampling on the lesser-understood ichneumonids, but also included sampling of braconids to enable comparison with a previously published 7-gene phylogeny on Braconidae (Sharanowski et al., 2011). We utilized 59 Ichneumonidae species representing 25 subfamilies and 21 Braconidae species representing 17 subfamilies (Table 1). For Ichneumonidae, taxa were selected to maximize the number of represented lineages and provide denser sampling for lineages that were suspected to be paraphyletic based on previous studies (Quicke et al., 2009) or contain PDVs (Bigot et al., 2008; Strand and Burke, 2015). For Braconidae, taxa were selected to include representatives of major lineages to assess how our results obtained with the anchored hybrid enrichment compare to a more traditional molecular study completed by Sharanowski et al. (2011). Species level identifications were not possible for the majority of taxa, as many genera require revisionary works or there are not adequate keys or descriptions of species to ensure accuracy. Outgroups consisted of genomic data from published model genomes that were included in our probe design (Table 1). We follow the higher-level groupings of Sharanowski et al. (2011) for Braconidae and Bennett et al. (2019) for Ichneumonidae, except we keep the grouping Labeniformes for members of Labeninae, and do not include this lineage within Ichneumoniformes *s*.*l*. All specimens were deposited in the University of Central Florida Collection of Arthropods (UCFC) except specimens from Sweden, which were deposited at Station Linné, Swedish Malaise Trap Project (SMTP).

### 2.3 Hybrid Enrichment Genome Capture & Data Processing

The remaining sequence data were obtained at Florida State University’s Center for Anchored Phylogenomics (www.anchoredphylogeny.com) using hybrid enrichment sequence capture with SureSelect. For this approach, genomic DNA samples were extracted using the Qiagen DNEasy kit with quality assessed using gel electrophoresis and a Nanodrop. Indexed libraries were prepared following a protocol modified from Meyer & Kircher (2010) but modified to accommodate the Beckmann-Coulter FXP liquid handling robot. Libraries were pooled and enriched using an Agilent Custom SureSelect kit containing all 57,066 probes and were sequenced using PE 100bp PE sequencing on the Illumina HiSeq 2000.

The resulting paired Illumina anchored hybrid enrichment reads were demultiplexed with no mismatches tolerated after quality filtering using the Illumina Casava high chastity filter. Read pairs were then merged using a Bayesian approach that identifies the sequence overlap, removes adapter sequences, and corrects sequencing errors in overlapping regions (Rokyta et al. 2012). The resulting reads were then assembled using a quasi de novo assembler described by Hamilton et al (2016), with the following references (Table 1): *Megachile rotundata*, *Microplitis demolitor*, *Aleiodes sp*. (PSUC_FEM 10030003), *Lissonota sp*. (PSUCFEM_10030012), and *Odontocolon* sp. (PSUC_FEM 10030002). The effects of low-level contamination and misindexing were avoided by filtering out assembly clusters with fewer than 100 reads. To establish orthology, homologous consensus sequences from each assembly cluster passing the coverage filter were evaluated to compute a pairwise distance matrix that was used to identify orthosets using a neighbor-joining approach (following Hamilton et al., 2016). After assembly and preliminary trimming using the pipeline, 479 loci were retained.

### 2.4 Alignment and Dataset Filtering

Orthologous sequences were aligned using MAFFT (v7.023b, with –genafpair and –maxiterate 1000 flags (Katoh and Standley, 2013a). Alignments were manually inspected for errors after autotrimming (mingoodsites=12, minpropsame=0.4 and 20 missing taxa allowed) following the approach outlined in Hamilton et al. (2016). Alignments were then manually checked to ensure the proper reading frame was retained after auto-trimming by using the gene maps of *Apis mellifera*, *Nasonia vitripennis*, and *Solenopsis invicta* as a guide, and where necessary, non-coding regions were removed (7 genes captured parts of introns and 3 genes captured parts of untranslated regions (UTRs).

### 2.5 Model testing and Phylogenetic Analyses

Each locus alignment was translated into a protein dataset and both protein and nucleotide datasets were concatenated using SequenceMatrix 1.8 (Vaidya et al., 2011) and analyzed separately. We analyzed the datasets using Maximum Likelihood with IQ-Tree v.1.6.10 (Nguyen et al., 2014) on either the Stokes HPC at the University of Central Florida Advanced Research Computing Center or on the CIPRES Science Gateway (Miller et al., 2009; Miller et al., 2010). Bayesian analyses were also performed on the concatenated nucleotide dataset with the third codon removed using MrBayes 3.2.6 (Ronquist et al., 2012). The full nucleotide or protein dataset was not completed using MrBayes, as analyses were taking longer than 3 months and often did not converge, even when constraints of known monophyletic lineages were applied. For the Nucleotide Bayesian analysis with the third codon position removed, we constrained monophyly for Braconidae, Ichneumonidae, Ants, and Bees. PartitionFinder V.2.1.1 (Lanfear et al., 2012; Lanfear et al., 2016) was used to find the best-fit models of molecular evolution and partitioning scheme using each locus as *a priori* subsets for the nucleotide dataset. For each treatment, an hcluster search with branch lengths linked was performed and model selection was based on the Bayesian Information Criterion (Schwarz, 1978). For the protein dataset, the best fit models of molecular evolution and partitioning scheme was inferred using ModelFinder (Chernomor et al., 2016; Kalyaanamoorthy et al., 2017) within IQ-Tree v.1.6.10, limited to the following models with the option for invariant and gamma parameters: Blosum62, Dayhoff, JTT, LG, VT, and WAG. All alignment datasets can be found on FigShare (10.6084/m9.figshare.12497468).

We also ran saturation, relative synonymous codon usage, and long branch exclusion tests and ran the nucleotide dataset with the 3^rd^ position removed using IQ-tree using the methods outlined above. Nodal support in IQ-tree analyses were assessed with 1000 pseudo-replicates of both the ultrafast bootstrap (Hoang et al., 2017) and Shimodaira-Hasegawa-like approximate Likelihood Ratio Test (SH-aLRT) (Guindon et al., 2010). The Bayesian analysis was run with 45 million generations sampling every 1000 generations with a 25% burn-in. The analysis consisted of two independent runs with four chains each. Convergence of the Bayesian analysis was assessed and confirmed using diagnostics in Tracer v.1.6 (Rambaut et al., 2013). To test for saturation, we calculated the number of pairwise differences and Tajima Nei genetic distance for each codon position using MEGA v.7.0 (Kumar et al., 2016), and plotted against each other using R (R Team, 2016) and the package “ggplot2”(Wickham, 2016). To test for codon bias in the nucleotide dataset we performed a principal component analysis using relative synonymous codon usage (RSCU) and amino acid composition, and performed a phylogenetic ANOVA using the effective number of codons (ENC) in R with the following packages: “seqinr” (Charif and Lobry, 2007), “phangorn” (Schliep, 2011), “geiger” (Harmon et al., 2008), “vhica” (Wallau et al., 2016), and “dplyr” (Wickham et al., 2015). In addition to this we calculated GC content for each taxon and synonymous skew on a scale of 0 to 1 for each of the six-fold degenerate amino acids (Rota-Stabelli et al., 2013) using the formula introduced by Perna and Kocher (1995), and mapped these metrics onto the nucleotide phylogeny using the R packages “ape” (Paradis et al., 2004) and “phytools” (Revell, 2012). As there were differences in some relationships between the nucleotide and amino acid analyses, we tested stability of the relationships recovered in the amino acid analyses by performing a series of taxon exclusion tests (Bergsten, 2005). For each analysis, one or more taxa were removed and the analysis rerun in IQ-tree following the methods outlined above. Choices of taxa to exclude were based on length of branches and instability in phylogenetic position between the nucleotide and amino acid datasets.

Successfully captured PDV genes were aligned along with the available GenBank sequences used to facilitate capture (FigShare 10.6084/m9.figshare.12497468) using Geneious with the native alignment algorithm and default parameters. These alignments were run through jmodeltest2 (Darriba et al., 2012) to assign models of evolution and were then run in MrBayes v.3.2.6 (10 M generations, 3 runs) using CIPRES Science Gateway (Miller et al., 2010). All were confirmed to have converged with Tracer v. 1.6 (Rambaut et al., 2013). As no taxa were captured with U1, these analyses included three genes: six taxa for *Bracovirus HzNVorf128* at 1035 aligned bases and run under a GTR+G model, four taxa for *Bracovirus per os infectivity factor p74* run with 2144 bps under a GTR+I model, and ten sequences for the *Ichnovirus IVSP4*, 1198 bp run under a GTR+I model.

### 2.6 Ancestral State Reconstructions

To examine evolutionary transitions of parasitoid biology, ancestral states were reconstructed for two traits: ectoparasitism (0) versus endoparasitism (1); and idiobiosis (0) versus koinobiosis (1) (Supplementary Table S1). Reconstructions were completed using the IQ-tree phylogeny of the amino acid analysis and repeated with Labeninae removed (due to instability in this lineage’s phylogenetic position; see results). To assess robustness of results across phylogenetic uncertainty, the analysis was repeated with the IQ-tree phylogeny based on nucleotide data with the third position removed (nt-3out) and repeated with Labeninae removed. Binary characters were analyzed using Maximum Likelihood using Mesquite version 3.2 (Maddison and Maddison, 2019). Two models were tested for likelihood reconstructions: the Markov k-state one parameter (Mk1), which has an equal rate for character gains and losses, and the asymmetrical two-parameter Markov k-state (Assym2), which applies a separate rate. To test whether the characters evolved in drastically different ways between Braconidae and Ichneumonidae, we tested whether the same model was a better fit when only one family was included for analysis.

## 3. Results

### 3.1 Sequence Data

Selective enrichment was good, with 27.3% of reads on average (range 14.6–49.0%) mapped to loci. Most of the loci were recovered across individuals, with an average of 478 of the 541 genes captured per individual. This is nearly the same number of genes that were recovered from blasts of taxa sequenced from whole genome data against references for each gene (n=480), suggesting that nearly all genes present with close affinity to reference sequences were captured. Many of the loci had comprehensive or nearly comprehensive coverage, with 206 of the loci having high read numbers (>100) mapped across all 35 taxa (1 taxon failed in the sequencing run) and 297 loci having high read mapping across at least 33 of the 35 taxa. Average length per locus was 990 bps (Supplementary Fig. 1). After trimming only seven genes had introns partially captured and three had portions of untranslated regions captured, which were manually trimmed out of the dataset. The concatenated datasets had a total of 91 taxa and 479 genes with lengths of 150,435 base pairs and 50,145 amino acids for the nucleotide and amino acid datasets, respectively. The nucleotide dataset with the third position removed had 100,290 base pairs.

### 3.2 Evolutionary Relationships among Ichneumonoidea

Saturation was indicated in the third codon position of the concatenated dataset (Fig. S2), and thus the maximum likelihood phylogeny with all data included (Fig. S3) is not discussed further given the likelihood for systematic error. The maximum likelihood amino acid phylogeny is depicted in Fig. 1A-C. Nodal supports for the nucleotide dataset with the 3^rd^ codon position removed (nt-3out) for the maximum likelihood (Fig. S4) and Bayesian (Fig. S5) phylogenies are also indicated on Fig. 1A-C where relationships were identical. All analyses recovered a maximally supported and monophyletic Ichneumonoidea, Braconidae, and Ichneumonidae (Fig 1a). Maximum support indicates a value of 100 for Shimodaira-Hasegawa approximate Likelihood Ratio Tests (aLRT), 100 for Ultra-Fast bootstraps (UFb), and 1.0 for posterior probability (PP).

**Figure 1.**
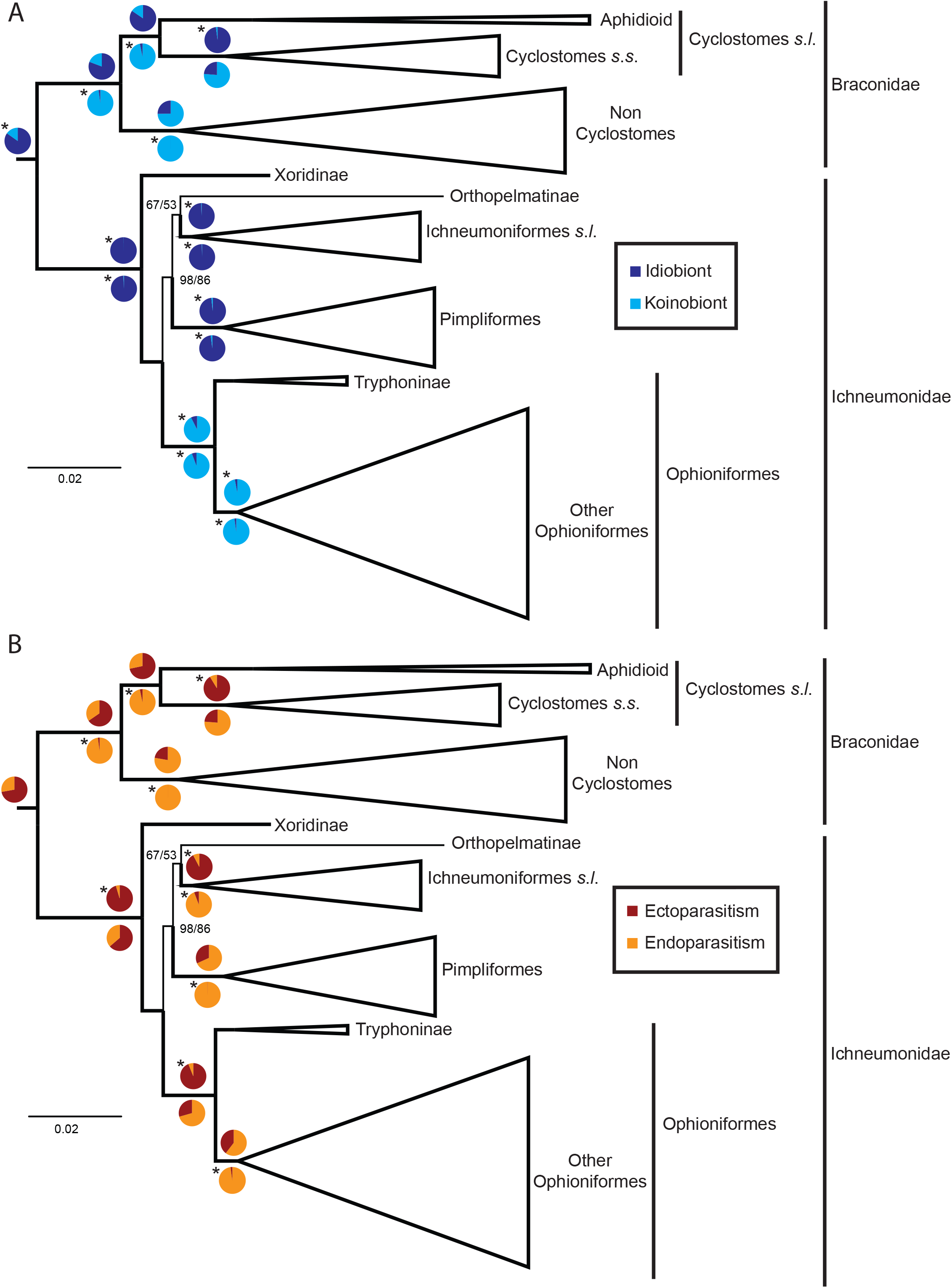
A-C. Maximum-likelihood (ML) phylogeny of Ichneumonoidea based on 479 genes analyzed as amino acids in IQ-tree. A) Summary tree with Braconidae and Ichneumonidae collapsed for viewing; B) Braconidae; C) Ichneumonidae. Branch supports are summarized in a 5 cell box, with the first two cells representing the amino acid analysis, the second two representing the ML phylogeny inferred from nucleotide data with the 3^rd^ position removed (nt-3out, see also Fig. S4), and the last box representing the Bayesian analysis of the nt-3out dataset (see also Fig. S5). For the ML analyses, two supports are given in the first and second boxes, respectively: the Shimodaira-Hasegawa approximate Likelihood Ratio Test statistic (aLRT) and the Ultra-Fast bootstrap value (UFb). The Bayesian posterior probability is reported in the last cell (PP). See branch support legend for color scheme of values for nodal supports. Taxa with an asterisk after the name have an uncertain identification at the lowest taxonomic rank indicated. The scale bar indicates the number of substitutions per site for branch lengths.

Within Braconidae, several higher-level lineages were also maximally supported across all analyses, including the cyclostomes *s*.*l*., cyclostomes *s*.*s*., non-cyclostomes, the Aphidioid, Helconoid and Microgastroid complexes, and the Alysioid subcomplex (Fig. 1B), similar to results reported in Sharanowski et al. (2011). Relationships among all cyclostomes were also consistent with Sharanowski et al. (2011), despite the low taxonomic sampling: the Aphidioid complex was sister to the cyclostomes *s*.*s*., Rhyssalinae was recovered as sister to the remaining cyclostomes *s*.*s*., and Braconinae was recovered as sister to the Alysioid complex with maximal support (Fig. 1B). The placements of Rogadinae and Pambolinae were labile between the amino acid (Fig. 1B) and nt-3out analyses (Figs. S4-5); however, support for Rogadinae as sister to Braconinae + the Alysioid complex increased when different outgroups were used (Fig. S6, Table S2) and when Aphidiinae was removed (Fig. S7). Many relationships within the non-cyclostomes were identical to results in Sharanowski et al. (2011). For example, Euphorinae was consistently recovered as sister to the Helconoid complex (Fig.1b): Acampsohelconinae (Brachistinae (Helconinae + Macrocentroid subcomplex)). One notable exception was the placement of Ichneutinae, which was variably placed as sister to Agathidinae + the Microgastroid complex (Fig. 1B) or sister to Agathidinae (Figs S4-5) with variable support. Most previous studies have recovered Ichneutinae as a paraphyletic group sister to the Microgastroid complex *s*.*s*. (Belshaw et al., 2000; Belshaw and Quicke, 2002; Dowton et al., 2002; Quicke and van Achterberg, 1990; Sharanowski et al., 2011). When Agathidinae was removed, Ichneutinae was recovered as sister to the Microgastroid complex (Fig. S8) with very high support (≥ 99), possibly indicating that low taxonomic sampling or long branch attraction may have been an issue for the unexpected placement of Ichneutinae.

Within Ichneumonidae, Xoridinae was always recovered as sister to all remaining ichneumonids with strong support (>95, Fig. 1C), consistent with previous analyses (Klopfstein et al., 2019; Quicke et al., 1999b) where Xoridinae was not set as the root (but see Bennett et al., 2019). The next basal lineage was Labeninae in the protein analysis (Fig. 1C), but this placement had very low support. In the nt-3out analyses, Labeninae was recovered as sister to Orthopelmatinae with relatively strong support (Figs. S4-S5), and these taxa were sister to Ichneumoniformes + Pimpliformes. Other recent studies with large taxon or gene sampling also had variable placement of Labeninae (Bennett et al., 2019; Klopfstein et al., 2019; Santos, 2017). These variable results may reflect the low taxonomic sampling, especially non-sampling of potentially related subfamilies, such as Brachycyrtinae, which has been recovered as the sister group to Labeninae (Bennett et al., 2019). In general, Labeninae was the most labile lineage across analyses and when different taxa were excluded (Table S2, Figs. S6-S17). When Xoridinae was excluded, the branch support for Labeninae as the most basal taxon is increased (Fig. S9). Further when Labeninae was excluded (Fig. S10), the support for all remaining Ichneumonids (except Xoridinae) as a monophyletic lineage was substantially increased (≥ 99).

Several higher-level relationships among Ichneumonidae were maximally supported, including Ichneumoniformes, Pimpliformes, and Ophioniformes (Fig. 1C). Ophioniformes was recovered as sister to all remaining ichneumonids (excluding Xoridinae and Labeninae) with the following relationships: Pimpliformes (Orthopelmatinae + Ichneumoniformes). The amino acid analysis recovered Orthopelmatinae as sister to the Ichneumoniformes, but with low support (Fig. 1C); however, in the outgroup test (Fig. S6) and several taxon exclusion tests, support for this relationship increased (Table S2, Figs. S6, S13-14, S17). Although Orthopelmatinae and Eucerotinae had very long branches, when Eucerotinae was excluded (Fig. S13), Orthopelmatinae was recovered as sister to Ichneumoniformes *s*.*l*. with increased support. Interestingly, when both Labeninae and Orthopelmatinae were excluded (Table S2, Fig. S12), support for the major backbone relationships increased (≥ 99): Ophioniformes (Ichneumoniformes + Pimpliformes).

Eucerotinae was always recovered as sister to a clade containing Adelognathinae, Cryptinae, and Ichneumoninae with maximal support (Fig. 1C). Despite our low taxonomic sampling for this lineage, these results support Eucerotinae within Ichneumoniformes *s*.*l*. (following Bennett et al., 2019). However, our lack of members of several subfamilies within Ichneumoniformes *s*.*l*. (Brachycyrtinae, Claseinae, Agriotypinae, Microleptinae, Pedunculinae, Ateleutinae, Phygadeuontinae) limits understanding of phylogenetic relationships in this lineage. The representatives we included for Ichneumoninae and Cryptinae were always recovered as monophyletic with maximum support, however, the placement of Adelognathinae varied with respect to these subfamilies across the amino-acid (Fig. 1C) and Bayesian nt-3out analysis (Fig. S5).

Similar to Klopfstein et al. (2019), relationships within Pimpliformes varied widely across the amino acid (Fig. 1C) analysis and nt-3out analyses (Figs. S4-5). In the amino acid analysis, Acaenitinae was recovered as sister to the remaining Pimpliformes with strong support (Fig. 1C), whereas Diplazontinae + Orthocentrinae were recovered at the base of the radiation in the nucleotide analyses with strong support (Figs S4-5). Interestingly, the relationships recovered in the amino acid analysis (Fig. 1C) remained stable across all outgroup and taxon exclusion tests (Table S2, Figs. S6-17) and were largely consistent with relationships proposed in a comprehensive morphological study by Wahl and Gauld (1998). We investigated codon use biases and observed that Diplazontinae has more similar codon usage to Ichneumoniformes *s*.*s*. than the rest of Pimpliformes (Fig. S18), and this may account for its more basal position in the nt-3out analyses (Figs. S4-5). Like long branch attraction (Bergsten, 2005), false homologies created through the codon use patterns may pull convergent taxa falsely together.

Consistent and well supported relationships within Pimpliformes included: Diplazontinae + Orthocentrinae (consistent with Wahl, 1990; Wahl and Gauld, 1998); Collyriinae + Cyllocerinae; and monophyly of the Ephialtini and Pimplini tribes of Pimplinae, consistent with the findings of Klopfstein et al. (2019) (Fig. 1C). Pimplinae was recovered as paraphyletic when nucleotide data was analyzed, but monophyletic when the amino acid data was analyzed (a similar issue was found in Klopfstein et al., 2019). There was also a distinct difference in codon usage between the Ephialtini and Pimplini clades of Pimplinae (Fig. S19). Thus, the amino acid analysis probably more accurately reflects the true monophyly of Pimplinae, which has been supported by some morphological studies (Gauld et al., 2002; Wahl and Gauld, 1998). The lone specimen of Poemeniinae (*Neoxorides* sp.) was recovered as sister to the Pimplinae, suggesting its close affiliation (Bennett et al., 2019; Quicke et al., 2009; Wahl and Gauld, 1998). The inclusion of Collyriinae in Pimpliformes supports previous studies (Bennett et al., 2019; Klopfstein et al., 2019; Quicke et al., 2009; Sheng et al., 2012).

Within Ophioniformes, the following subfamilies (that had more than one exemplar) were recovered as monophyletic across all analyses with maximum support (Fig. 1C): Tryphoninae, Banchinae, Tersilochinae, Metopiinae, Anomaloninae, Cremastinae, and Campopleginae. Tryphoninae was consistently recovered and maximally supported as the sister to all remaining Ophioniformes. Banchinae was the next most basal lineage (consistent with Klopfstein et al., 2019; Quicke et al., 2009), but this placement was poorly supported (Fig. 1C). However, this placement was largely consistent across the outgroup and taxon exclusion analyses (Table S2), except when Mesochorinae was removed (Fig. S16). Further, in the nt-3out analyses (Figs S4-5), Banchinae was recovered as sister to Tersilochinae with strong support, and both were sister to the remaining Ophioniformes (except Tryphoninae) (Figs. S4-5). The remaining Ophioniformes were consistently recovered in a maximally supported clade, including: Mesochorinae, Oxytorinae, Ctenopelmatinae, Metopiinae, Cremastinae, Hybrizontinae, Anomaloninae, Ophioninae, and Campopleginae. Of these, Mesochorinae was recovered as sister to the rest, but with variable support (Fig. 1C). Ctenopelmatinae was unsurprisingly paraphyletic as was found previously (Bennett et al., 2019; Quicke et al., 2009). Metopiinae was consistently recovered as sister to a maximally supported clade that roughly corresponds to Quicke et al.’s (2009) “higher Ophioniformes,” including Cremastinae, Hybrizontinae, Anomaloninae, Ophioninae, and Campopleginae. Interestingly, Hybrizontinae (a group that has varied widely in placement across all previous analyses) was recovered as sister to Cremastinae with strong but variable support across all analyses (Fig. 1C). When Hybrizontinae was excluded (Fig. S15), Anomaloninae was recovered as sister to Cremastinae with relatively high support (>93). Ophioninae was consistently recovered as the sister group to Campopleginae with maximal support across all analyses.

### 3.3 Ancestral State Reconstructions

Ancestral biological states were reconstructed on phylogenies that varied in the data type they were estimated from (amino acid or nt-3out) and taxon representation: either all taxa were included, all taxa but Labeninae (due to its uncertain placement), and with or without outgroups included. Further, each family was tested individually based on the amino acid analysis, with all other taxa removed. As our sampling was not comprehensive, we only discuss ancestral state reconstructions for higher-level relationships, including Ichneumonoidea, Braconidae and Ichneumonidae. Within Braconidae we also discuss reconstructions for the ancestors of the Cyclostomes *s*.*l*., Cyclostomes *s*.*s*., and Non-cyclostomes, and within Ichneumonidae, ancestors of Ichneumoniformes *s*.*l*., Pimpliformes, Ophioniformes, and Ophioniformes excluding Tryphoninae (Fig. 2A and B). The full reconstructions for each trait and input phylogeny can be found on FigShare (10.6084/m9.figshare.12497468).

**Figure 2.**
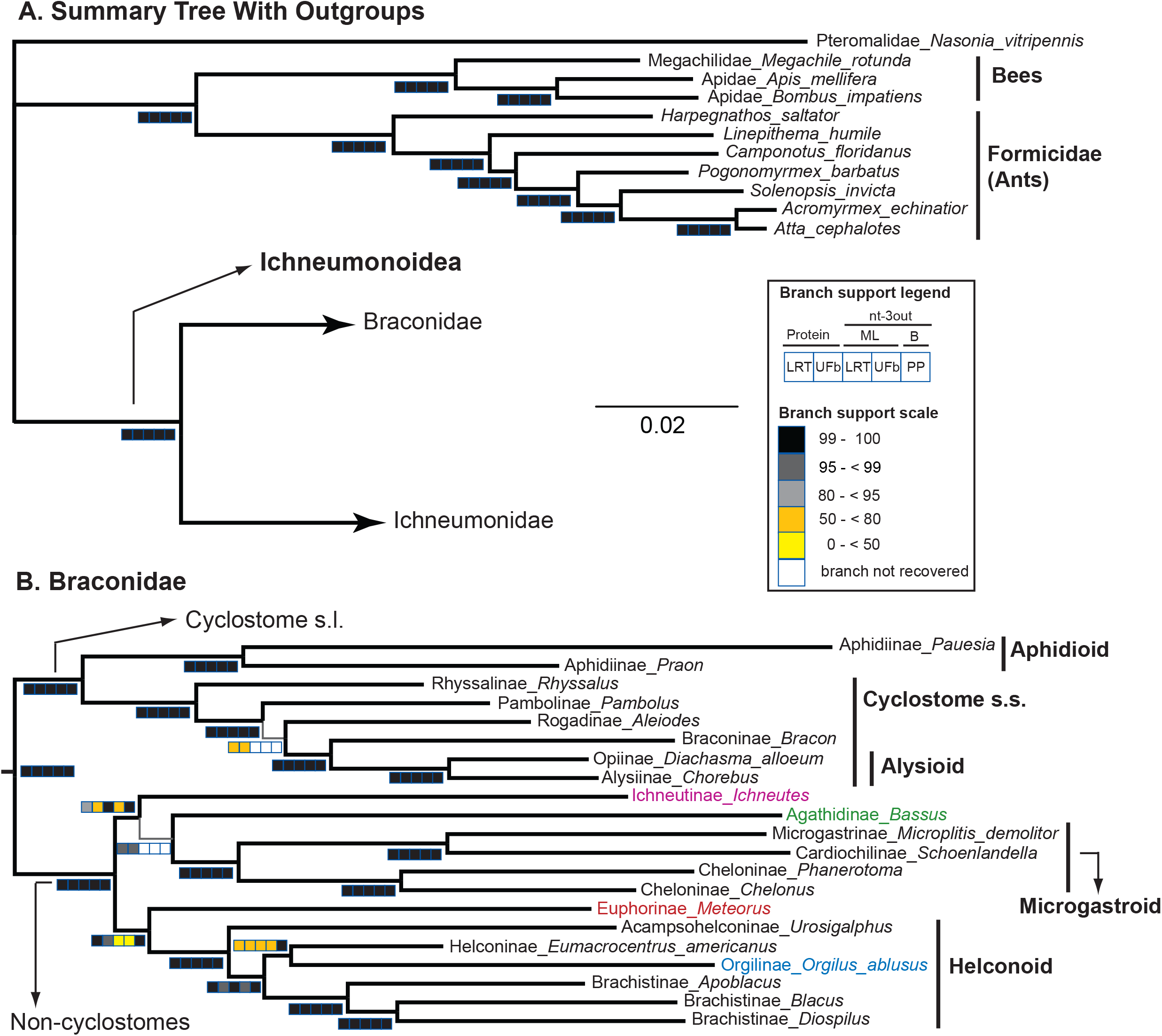
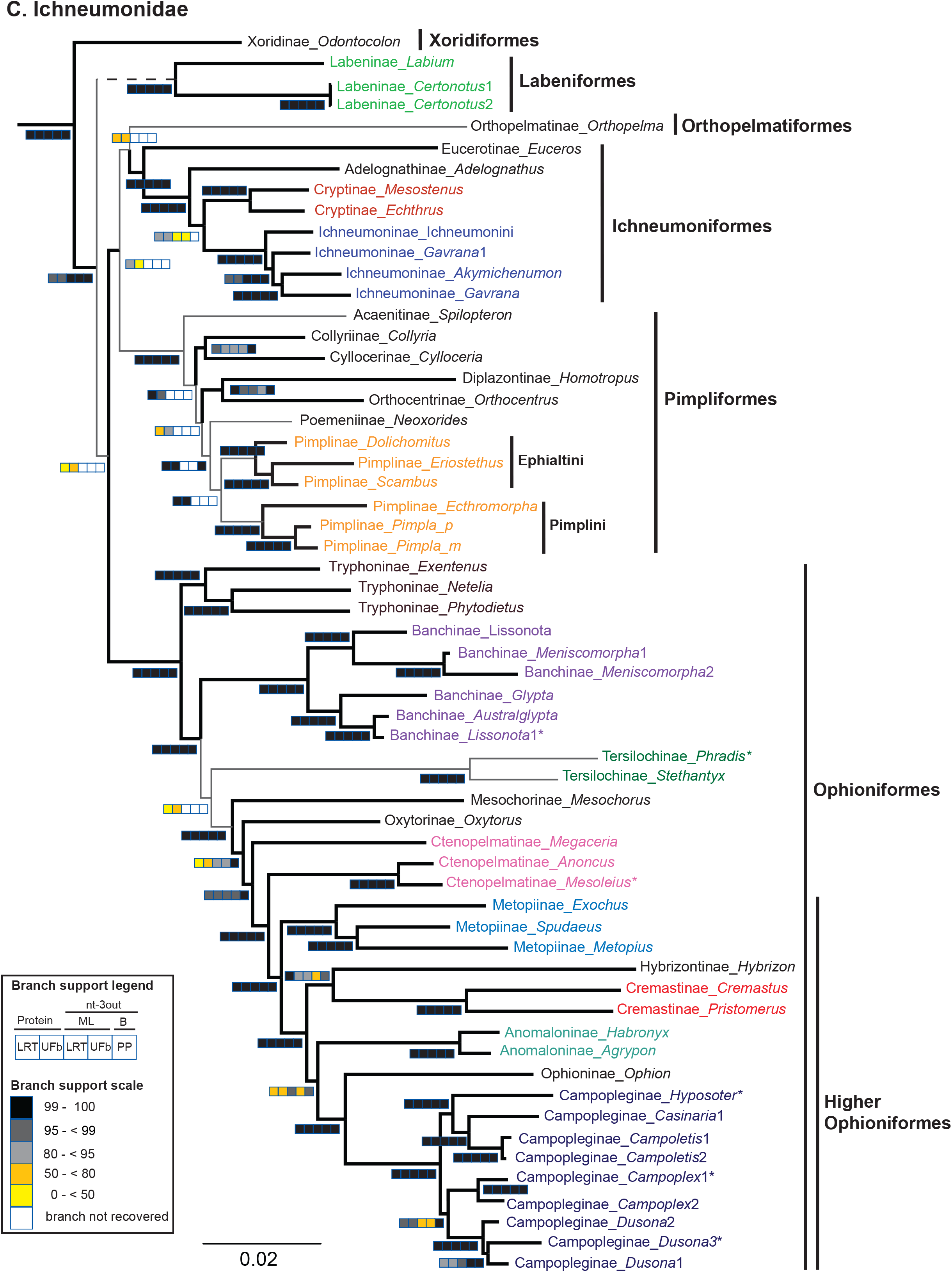
A-B. Summary of two ancestral state reconstructions (ASR) using maximum likelihood for A) Idiobiosis versus koinobiosis and B) Ecto- versus endoparasitism. The collapsed tree visualized here is based on the maximum likelihood phylogeny of amino acid data with Labeninae removed (Fig. S10). Pie charts show the proportional likelihood (PLH) for each character state and an asterisk near the chart indicates the reconstruction as significant for the state with the highest PLH (see also Table 3). Pie charts above the nodes have all taxa included, whereas pie charts below the nodes were ASRs run with all taxa removed except the family of interest for both Braconidae and Ichneumonidae.

The Assym2 model (separate parameters for gains and reversals) consistently outperformed the Mk1 model (equal rates of transition between character states) for character 1 (idiobiont vs koinobiont) when all Ichneumonoidea were analyzed together (Table 2). Under these analyses, the ancestors of Ichneumonoidea, Ichneumonidae, and Braconidae were reconstructed as idiobionts, although this was only consistently significant for Ichneumonidae (Fig. 2A, Table 3). Within Braconidae, the ancestors of the Cyclostomes *s*.*l*. and *s*.*s*. were also recovered as idiobionts, but this was only consistently significant for the latter. Unsurprisingly, the ancestor of the non-cyclostomes was significantly koinobiont as all known members are koinobiont (Fig. 2A, Table 3). Within Ichneumonidae, the Ichneumoniformes *s*.*l*. and Pimpliformes were ancestrally idiobiont, whereas Ophioniformes was koinobiont, and all reconstructions were significant (Fig. 2A, Table 3). When the families were analyzed individually, the above results remained consistent for Ichneumonidae, but not Braconidae as the Mk1 model was preferred for Braconidae (Table 2). Under this model all major braconid nodes were reconstructed as koinobiont and significant for all lineages except the cyclostomes *s*.*s*. (Fig. 2A, Table 3), suggesting this character is sensitive to the data used to infer the ancestral states for Braconidae.

**Table 2.** Model tests for maximum likelihood ancestral state reconstructions for character 1 (idiobiosis vs. koinobiosis) and character 2 (ecto- vs. endoparasitism). Two models were tested, the Markov k-state one parameter (Mk1), which has an equal rate for character gains and losses, and the asymmetrical two-parameter Markov k-state (Assym2), which applies a separate rate to gains and losses. Likelihood ratio tests were performed to assess which model better fit the data, which are chi-square (χ2) distributed and critical values was used to assess significance at p<0.01. The best supported model for each source phylogeny is highlighted in gray. Source phylogenies include the amino acid maximum likelihood phylogeny (AA w/ OGs) with all taxa (Fig 1. A-C); and with outgroups (OGs) trimmed prior to analysis (AA w/o OGs); the amino acid phylogeny with Labeninae excluded (AA w/ OGs Lab-out, Fig. S10); and with outgroups trimmed (AA w/o OGs Lab-out); the nucleotide dataset with the third position removed (nt-3out, Fig. S4). The other three source phylogenies are based on the amino acid dataset (Fig. 1A-C), either with (AA Bracs only; AA Ichs only) or without Labeninae excluded (Fig. S10; AA Ichs only Lab-out), and all taxa were trimmed except the family of interest: Braconidae (Bracs) or Ichneumonidae (Ichs).

| Source Phylogeny | Character | Log Likelihood | | $\chi^2$ | P-value |
| --- | --- | --- | --- | --- | --- |
|  |  | Mk1 | Assym2 |  |  |
| AA w/ OGs | 1 | -34.216 | -27.230 | 13.973 | 0.00019 |
|  | 2 | -33.670 | -29.275 | 8.790 | 0.003 |
| AA w/o OGs | 1 | -33.267 | -25.052 | 16.430 | 0.00005 |
|  | 2 | -33.180 | -29.142 | 8.076 | 0.0045 |
| AA w/ OGs Lab-out | 1 | -31.663 | -25.924 | 11.478 | 0.0007 |
|  | 2 | -30.970 | -27.608 | 6.724 | 0.0095 |
| AA w/o OGs Lab-out | 1 | -30.556 | -23.539 | 14.033 | 0.00018 |
|  | 2 | -30.585 | -27.501 | 6.169 | 0.013 |
| nt-3out | 1 | -35.345 | -27.692 | 15.307 | 0.000092 |
|  | 2 | -30.625 | -33.386 | -5.522 | 0.019 |
| AA Bracs only | 1 | -10.262 | -8.207 | 4.110 | 0.043 |
|  | 2 | -10.262 | -8.207 | 4.110 | 0.043 |
| AA Ichs only | 1 | -22.721 | -17.510 | 10.423 | 0.0012 |
|  | 2 | -22.571 | -21.315 | 2.511 | 0.119 |
| AA Ichs only Lab-out | 1 | -20.590 | -15.865 | 9.450 | 0.0021 |
|  | 2 | -20.594 | -19.594 | 1.999 | 0.16 |

**Table 3.** Summary of ancestral state reconstructions across several source phylogenies for character 1 (idiobiosis vs. koinobiosis) and character 2 (ecto- vs. endoparasitism). Color coding and the proportional likelihoods (PLH) are listed for the most likely character state. Source phylogenies are detailed in Table 2.

| Node | AA | AA (no OGs) | AA (Lab-out) | AA (no OGs, Lab-out) | nt-3out | AA Bracs only | AA lchs only | AA lchs only (Lab-out) |
| --- | --- | --- | --- | --- | --- | --- | --- | --- |
|  | PLH | PLH | PLH | PLH | PLH | PLH | PLH | PLH |
| Ichneumonoidea | **0.941** | 0.877 | **0.899** | 0.848 | **0.928** | N/A | N/A | N/A |
| Braconidae | 0.871 | 0.832 | 0.828 | 0.807 | 0.826 | **0.979** | N/A | N/A |
| Cyclostomes *s.l.* | **0.892** | 0.864 | 0.857 | 0.845 | 0.849 | **0.968** | N/A | N/A |
| Cyclostomes *s.s.* | **0.986** | **0.986** | **0.976** | **0.98** | **0.974** | **0.762** | N/A | N/A |
| Non-cyclostomes | **0.815** | **0.784** | **0.777** | **0.748** | **0.836** | **0.999** | N/A | N/A |
| Ichneumonidae | **0.999** | **0.999** | **0.989** | **0.999** | **0.996** | N/A | **0.96** | **0.994** |
| Ichneumoniformes *s.l.* | **0.997** | **0.998** | **0.991** | **0.994** | **0.998** | N/A | **0.983** | **0.996** |
| Pimpliformes | **0.98** | **0.986** | **0.974** | **0.983** | **0.997** | N/A | **0.971** | **0.981** |
| Ophioniformes | **0.956** | **0.945** | **0.939** | **0.925** | **0.96** | N/A | **0.912** | **0.943** |
| Ophioniformes (no Try) | **0.986** | **0.98** | **0.978** | **0.97** | **0.99** | N/A | **0.96** | **0.979** |
| Ichneumonoidea | 0.788 | 0.794 | 0.724 | **0.987** | 0.861 | N/A | N/A | N/A |
| Braconidae | 0.714 | 0.715 | 0.66 | **0.995** | **0.96** | **0.979** | N/A | N/A |
| Cyclostomes *s.l.* | 0.763 | 0.762 | 0.717 | **0.979** | **0.94** | **0.968** | N/A | N/A |
| Cyclostomes *s.s.* | **0.945** | **0.943** | **0.918** | 0.768 | 0.732 | 0.762 | N/A | N/A |
| Non-cyclostomes | **0.805** | **0.81** | 0.772 | **0.999** | **0.999** | **0.999** | N/A | N/A |
| Ichneumonidae | **0.994** | **0.994** | **0.959** | **0.97** | 0.84 | N/A | 0.807 | 0.637 |
| Ichneumoniformes *s.l.* | **0.965** | **0.961** | **0.932** | **0.998** | **0.958** | N/A | **0.989** | **0.942** |
| Pimpliformes | 0.745 | 0.762 | 0.664 | **0.999** | **0.999** | N/A | **0.999** | **0.998** |
| Ophioniformes | **0.97** | **0.968** | **0.945** | **0.958** | 0.884 | N/A | **0.906** | 0.706 |
| Ophioniformes (no Try) | 0.647 | 0.655 | 0.598 | **0.998** | **0.995** | N/A | **0.994** | **0.979** |

For character 2 (ecto- vs. endoparasitism), the ancestral states were very sensitive to the phylogeny utilized in the analysis (Fig. 2B). The Mk1 model was supported for character 2 when Labeninae and outgroups were both excluded, when the nt-3out phylogeny was used, or when the Braconidae or Ichneumonidae were analyzed individually (Table 2). Under the Mk1 model, all major nodes were reconstructed as endoparasitoid and this consistently significant for most reconstructions except the Ichneumonoidea, Ichneumonidae, Ophioniformes, and cyclostomes *s*.*s*. (Fig. 2B, Table 3). For Ichneumonidae, these reconstructions suggested that the Ichneumonidae, Ichneumoniformes *s*.*l*., and Pimpliformes had idiobiont endoparasitoid ancestors. When the Assym2 model was most appropriate (Table 2), all major braconid nodes were reconstructed as ectoparasitoid, except the non-cyclostomes, and only the cyclostomes *s*.*s*. had significant reconstructions (Table 3). There was also inconsistency across the Ichneumonidae, where under the Assym2 model, only the Pimpliformes and Ophioniformes without Tryphoninae were recovered as endoparasitoid and these reconstructions were not significant (Fig. 2B, Table 3). These data suggest that ancestral reconstruction, particularly of ecto- vs. endoparasitism, is sensitive to the taxon representation and type of data used in the analysis.

### 3.4 Capture of Polydnavirus Genes

Capture of polydnavirus genes had limited success. The *U1* ichnovirus replication gene was not captured in any of the taxa in our dataset including the campoplegines most likely to harbor a copy of it. The IVSP4 gene was represented in the probe set by two paralogs from *Hyposoter didymator* (IVSP4-1 and IVSP4-2) and one gene copy from *Campoletis sonorensis* (Fig. 3A). *Glypta fumiferanae* (Banchinae) was not used as the probes were designed prior to the publication of this genome (Béliveau et al., 2015). We were able to capture matches to both IVSP4-1 and IVSP4-2 in another *Hyposoter* species with considerable divergence from the other *Hyposoter* (I9995), suggesting these genes may be rapidly evolving. We were also able to capture several copies of this gene family from *Campoletis*: two individuals of *Campoletis* (I2339, I10007, Table 1) matched very closely to the original *Campoletis sonorensis* sequence and *Campoletis* sp. 1 (I10007) exhibited two additional paralogs (copy 2, 3) that were more closely related to the other *Campoletis sp. 1* paralog than to paralogs of *Hyposoter* (Fig. 3A). *Casinaria* sp. showed affinities to a copy in *Campoletis* (copy 3). This demonstrates that this ichnovirus replication gene is part of a gene family that likely has lineage specific diversification. *Hyposoter, Campoletis* sp. 1 and 2, and *Casinaria* sp. 1 are recovered in a maximally supported lineage and represent just one lineage of Campopleginae (Fig. 1C). Interestingly, these genes were not captured in species of *Dusona* and *Campoplex*, suggesting either these genera do not have ichnoviruses, or the ichnovirus genes are highly divergent from other lineages within Campopleginae. No IVSP4 family members were captured in *Lissonota* despite being present in the genome (Burke et al., 2020), but this may be due to not having a banchine sequence in the original alignment used to design the probes. Although in *Hyposoter* both IVSP4 copies were captured, it is interesting that not all copies were captured in each case, which again could be a complication of this approach for capturing gene copies successfully or for indication of potential gains and losses of ancestrally attained gene copies.

**Figure 3.**
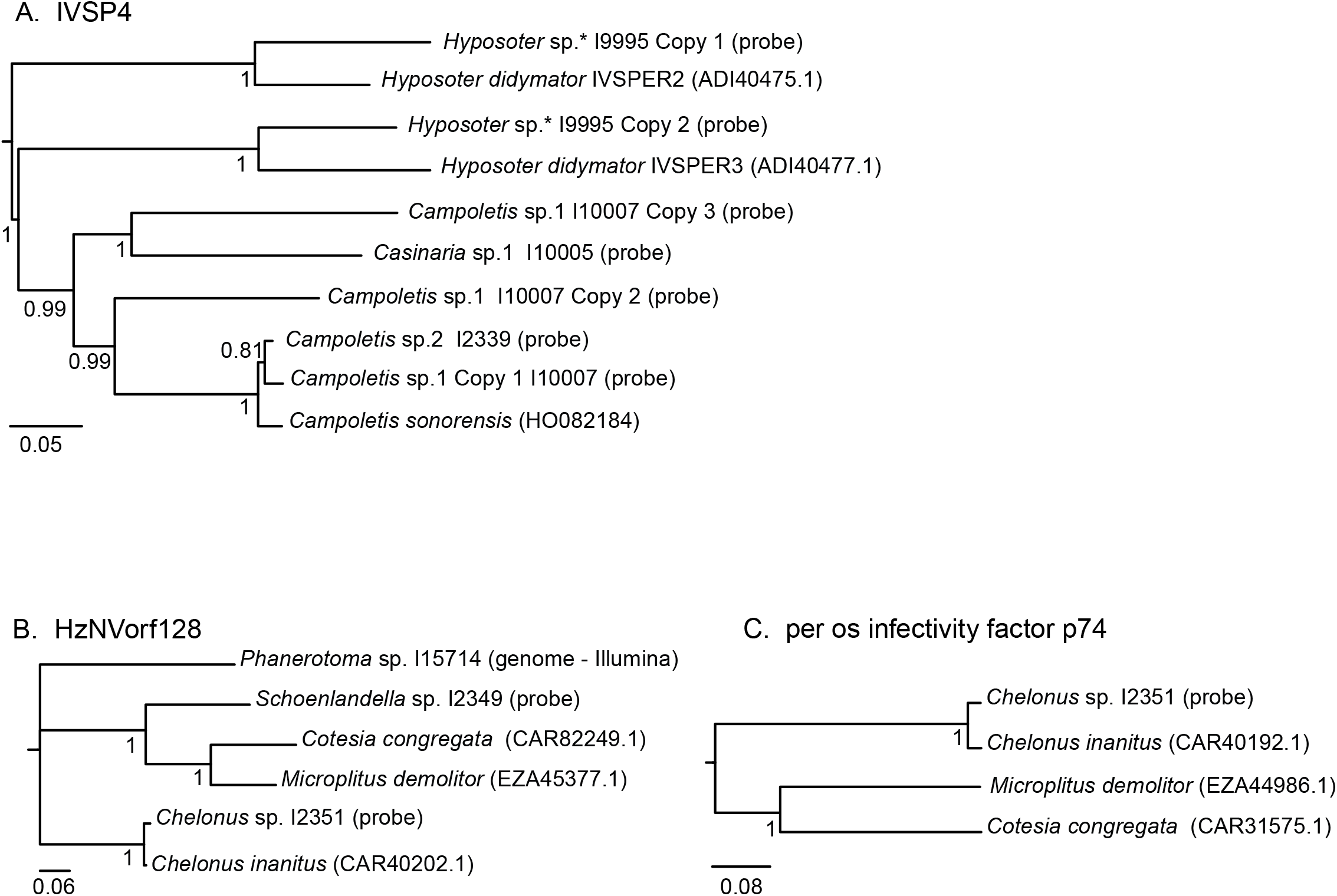
A-C. Bayesian phylogenies of three polydnavirus (PDV) genes: A) IVSP4, B) HzNVorf128, and C) per os infectivity factor p74. Sequences were either captured with anchored hybrid enrichment probes (probe), obtained from newly sequenced genomes (genome-Illumina), or were used as original reference sequences for probe capture and are available on GenBank (GenBank ID provided). Taxon voucher codes are listed next to genus name for newly sequenced taxa (FSU code, Table 1).

In the bracovirus replication genes, *HzNVorf128* was represented by a *Chelonus* (Cheloninae) and a *Cotesia* (Microgastrinae) species in the original probe design and was effectively captured and located for all taxa in our sampling thought to include bracoviruses (Fig. 3B). The relationships inferred from *HzNVorf128* were congruent with the expected phylogenetic relationships (Fig. 1B), suggesting the expected co-phylogeny of the replication machinery with the taxa they harbor (Fig. 3B). Also, there was no indication of gene duplications in this gene, making it a good candidate to evaluate further in PDV analysis. The *per os infectivity factor p74*, however, was captured/identified in the genomes of the two closely related microgastroids to the probe sequences, with *Schoenlandella* (Cardiochilinae) not captured and the gene not found in the *Phanerotoma* genomic sequence (Fig. 3C). It likely is evolving more rapidly, thus eluding successful identification beyond close relatives. We included the vankyrin genes to test if we could delineate members of a gene family using hybrid capture and because they are present in each of the lineages that have PDVs (bracoviruses and ichnoviruses). Some individuals exhibited an overabundance of vankyrins in the capture, e.g., 2% of the *Australoglypta* (Banchinae) sequence was comprised solely of vankyrins, and other individuals had very low and likely non-specific capture. Given the complexity of these results and establishing paralogy among divergent gene family members, we excluded vankyrins from further analysis.

## 4. Discussion

### 4.1 Utility of Probe Set for Phylogenetics

The designed 541 probes set worked very well for capturing genes across Ichneumonoidea, with an average of 478 genes captured per individual and 491 genes used in the final dataset analyzed here. The probes were designed from alignments across taxa within the Holometabola and thus genes were largely conserved. Further, the vast majority of genes captured only coding regions after trimming, making them appropriate genes for examining macroevolutionary relationships, such as for Ichneumonoidea, which has been estimated to have diverged in either the late Jurassic or early Triassic (Peters et al., 2017; Zhang et al., 2015). These probes have also been successfully used to capture loci for examining phylogenetic relationships and the evolution of eusociality in Vespidae (Piekarski et al., 2018), indicating they have wide applicability across different hymenopteran taxa.

### 4.2 Evolutionary Relationships among Ichneumonoidea

Sampling across the superfamily was sparse relative to the total number of described species within Ichneumonoidea (> 44,000, Yu et al., 2012), however, the inferred phylogenies were largely stable. This was true across different analytical methods and datasets and when taxa with suspected long branches or labile placement were excluded. Numerous maximally supported higher-level relationships were inferred using the 491 gene dataset, providing a robust higher-level evolutionary picture for Ichneumonoidea.

Although the taxon sampling for Braconidae was very low, the core higher-level relationships recovered from traditional molecular phylogenetic approaches with a handful of genes (Sharanowski et al., 2011) are largely recapitulated using a phylogenomics approach with denser genetic sampling. Interestingly, regions of the Braconidae phylogeny with low support were similar to regions of low support from Sharanowski et al. (2011), a symptom typical of rapid radiations (Giarla and Esselstyn, 2015; Xi et al., 2012), including in Braconidae (Banks and Whitfield, 2006; Zaldivar-Riverón et al., 2008). Thus, a combined dense taxonomic and genetic sampling approach will be necessary to fully understand the phylogenetic history of this extremely diverse family of parasitoid wasps.

The relationships of Ichneumonidae were also largely stable and highly supported with a few exceptions. All analyses and datasets recovered Xoridinae as the sister to all remaining Ichneumonids, an important finding since some previous analyses have used Xoridinae to root the Ichneumonid phylogeny (Quicke et al., 2000; Quicke et al., 2009) and its placement impacts the understanding of the evolution of parasitism (Belshaw and Quicke, 2002; Gauld, 1988). Subfamilies with previously ambiguous placement include Orthopelmatinae and Eucerotinae (Quicke et al., 2000; Quicke et al., 2009). In our analyses, Orthopelmatinae was recovered as sister to Ichneumoniformes *s*.*l*. with low support in the amino acid analysis. However, support for this relationship increased when different outgroups were used or when specific taxa were individually excluded (e.g. Eucerotinae, Adelognathinae). This relationship was not recovered in the nt-3out relationship because of the placement of Labeninae, but still suggested an affinity with Ichneumoniformes *s*.*l*. Orthopelmatinae has been considered an aberrant group, sometimes placed in its own Orthopelmatiformes group (Quicke et al., 2000; Quicke et al., 2009), of which the placement has been unstable across numerous previous analyses, either as sister to Ophioniformes (Quicke et al., 2000; Quicke et al., 2009), sister to Pimpliformes (Quicke et al., 2000, morphological analysis), as the second most basal lineage next to Xoridinae (under parsimony, Bennett et al., 2019), or within the Ophioniformes (Quicke et al., 2009). Whether or not Orthopelmatinae is sister to Ichneumoniformes *s*.*l*. will require denser taxonomic sampling, specifically across more basal lineages.

Eucerotinae has also been variably treated due to its unique (within Ichneumonoidea) larval biology and stalked eggs (see Bennett et al., 2019 for detailed review), but has more recently been associated with taxa not sampled here, including Agriotypinae, Claseinae, Pedunculinae, Brachycyrtinae, and/or Labeninae, along with Ichneumoniformes (Bennett et al., 2019; Gauld and Wahl, 2002; Quicke et al., 2009). Here, Eucerotinae was consistently recovered as sister to Ichneumoniformes with maximum support, consistent with recent studies with more extensive taxonomic sampling (Bennett et al., 2019; Santos, 2017). Further our data supports the assertion of Bennett et al. (2019) that Eucerotinae should be placed within the Ichneumoniformes (Bennett et al.’s Ichneumoniformes *s*.*l*.).

While monophyletic, the placement of Labeninae varied widely across analyses and taxon exclusion schemes, and this impacted the support values of a handful of deeper nodes within Ichneumonidae. When labile taxa were removed or different outgroups were used, support values tended to increase across the backbone of the ichneumonid phylogeny. We recovered Labeninae as sister to the rest of the Ichneumonidae, except Xoridinae, similar to some analyses in other previous studies (Klopfstein et al., 2019; Quicke et al., 2009; Santos, 2017). The placement of Labeninae has varied widely across these and other studies (Bennett et al., 2019; Quicke et al., 2000) and its true placement represents an important future challenge for understanding ichneumonid evolution.

Quicke et al. (2009) separated Ophioniformes into the ‘higher Ophioniformes’ (Campopleginae, Cremastinae, Anomaloninae, Ophioninae, Mesochorinae, Nesomesochorinae, and Tatogastrinae, the last two not sampled here) and a ‘basal grade’ of other subfamilies (Tryphoninae, Banchinae, Tersilochinae, Oxytorinae, Ctenopelmatinae, Metopiinae, Neorhacodinae, Sisyrostolinae, and Stilbopinae, the last three not sampled here). Alternatively, Bennett et al. (2019) defined the ‘higher Ophioniformes’ as including only Anomaloninae, Campopleginae, Cremastinae, Ophioninae, and Nesomesochorinae, but did not state why. Perhaps the smaller set of taxa relates to support for this grouping across some of their analyses, or maybe due to the somewhat contradictory statement of the groups’ membership in the discussion of Quicke et al. (2009): “The probable monophyly of the “higher” ophioniformes (Anomaloninae, Campopleginae, Cremastinae, Nesomesochorinae, Ophioninae, but not Tersilochinae nor Tatogastrinae) implies a very large ichneumonid radiation associated with koinobiont endoparasitism of Lepidoptera larvae (with some secondary host shifts within these groups). The hosts of Nesomesochorinae are completely unknown and the Hybrizontinae may or may not be associated with this clade (p. 1355)”. Further, in some of their analyses and other studies (Belshaw and Quicke, 2002; Quicke et al., 2000) Metopiinae is recovered with several of these subfamilies. Here we recovered a maximally supported group including Metopiinae, Hybrizontinae, Cremastinae, Anomaloninae, Ophioninae, and Campopleginae. The latter six subfamilies were also maximally supported across all analyses.

The placement of Hybrizontinae has varied widely across different analyses, including with affiliation to Ctenopelmatinae (Bennett et al., 2019), Lycorininae, Anomaloninae, and Ophioninae (Quicke et al., 2009) among others (see Bennett et al., 2019; Quicke, 2015 for a detailed review). Here, Hybrizontinae is consistently (albeit variably supported) as sister to Cremastinae and falls firmly within the ‘higher Ophioniformes’, contrary to (Bennett et al., 2019). Ophioninae was consistently recovered as the sister group to Campopleginae with maximal support across all analyses, a relationship espoused by Gauld (1985) and also recovered by Klopfstein et al. (2019). This finding is contrary to Bennett et al. (2019) and Wahl (1991) who recovered Cremastinae as sister to Campopleginae. Although the study of Klopfstein et al. (2019) was focused on Pimpliformes, they recovered very similar relationships among the Ophioniformes using an independent probe set.

There were some differences between the two datasets (amino acid and nt-3out), particularly for Ichneumonidae. However, several of these differences could be explained by further analysis of the data. For example, the relationships among Pimpliformes were completely different between the protein and nucleotide datasets, as was found in a recent phylogenomics study of this lineage (Klopfstein et al., 2019). Examination of codon biases showed that there were distinct differences across several lineages. Diplazontinae has a codon use pattern very similar to Ichneumoniformes, and thus, this lineage may have been pulled to the base of the Pimpliformes in the nt-3out analyses due to convergence. Similarly, Pimplinae was only recovered as monophyletic with the amino acid analysis due to distinct codon use patterns between the two tribes sampled here: Ephialtini and Pimplini (Delomeristini and Theroniini were not sampled). Pimplinae has been recovered as monophyletic with morphological data (Gauld et al., 2002; Wahl and Gauld, 1998), but paraphyletic with molecular data (Bennett et al., 2019; Klopfstein et al., 2019; Quicke et al., 2009). Interestingly, our amino acid results of Pimpliformes are most consistent with the relationships proposed by Wahl and Gauld (1998), but as mentioned by Klopfstein et al. (2019), the evolution of this lineage is likely the result of a rapid radiation which may require extensive genetic and taxonomic sampling to resolve.

Although nucleotide bias has been observed before between Braconidae and Ichneumonidae in select genes, mostly mitochondrial (Li et al., 2016; Quicke et al., 2020b; Sharanowski et al., 2010), here we note distinct codon use patterns across hundreds of genes. Further, the outgroups used here were most similar to Braconidae, suggesting that careful examination of codon biases, outgroup choice, and performing differential outgroup analyses are important for phylogenetic inference of Ichneumonoidea. Thus, it may not be appropriate to root braconid or ichneumonid phylogenies with its sister family. We also suggest that amino acid analyses are more useful for examining relationships among Ichneumonoidea, and particularly Ichneumonidae because of codon use patterns.

### 4.3 Evolution of Modes of Parasitism

Although the taxon sampling is low in this study, we did recover interesting patterns when inferring the ancestral states for major lineages in Ichneumonoidea. In general, it appears that the ancestor of Ichneumonoidea was an idiobiont ectoparasitoid, although the result was only occasionally significant for idiobiosis and never for ectoparasitism. Within Braconidae, the results varied depending on the data used or taxa included in the analyses. The ancestor of Braconidae was inferred as either an idiobiont ectoparasitoid or a koinobiont endoparasitoid. However, only the latter inferred biology was significant, a result that is consistent with Sharanowski (2009), which had denser taxonomic sampling for Braconidae. Koinobiont endoparasitism is the most common biology within Braconidae (Gauld, 1988), as this biology is common to several subfamilies within the cyclostomes *s*.*l*. and all of the non-cyclostomes. However, the non-cyclostomes were always recovered with a koinobiont endoparasitoid ancestor, whereas the cyclostomes *s*.*s* had the most support for an idiobiont ectoparasitoid ancestor, although the result varied across analyses. If the ancestor of Braconidae was a koinobiont endoparasitoid, then there may have been one or more transitions to idiobiosis and ectoparasitism within the cyclostomes *s*.*s*.

Within Ichneumonidae, the results for idiobiosis/koinobiosis were consistent across all analyses. Ichneumonidae, Ichneumoniformes *s.l.*, and Pimpliformes were all significantly reconstructed as idiobionts, whereas the ancestors for Ophioniformes, with or without Tryphoninae, were significantly reconstructed as koinobionts. As all Ophioniformes are koinobionts, this makes intuitive sense and demonstrates that koinobiosis developed early in ichneumonid evolution. For the ecto- versus endoparasitism, there was a great deal of inconsistency across the analyses. The only consistent results were for ancestors of Pimpliformes and Ophioniformes (excluding Tryphoninae), which were always reconstructed as endoparasitoids with varying statistical support. This suggests that Pimpliformes had an idiobiont endoparasitoid ancestor. Gauld (1988) theorized that idiobiont parasitoids transitioned from ectoparasitism to endoparasitism by attacking cocooned hosts, allowing for enough time to develop before the host metamorphoses into an adult. However, previous analyses (Klopfstein et al., 2019) have been equivocal with respect to the ancestral state of idiobiosis or koinobiosis for Pimpliformes, and the diversity of biological strategies in this group may make it one of the most interesting lineages to examine to understand the evolution of parasitism. The ancestor of all Ophioniformes may have had a koinobiont ecto- or endoparasitoid ancestor, as results varied depending on analysis. As Tryphoninae are koinobiont ectoparasitoids, this may have been a transitory step from idiobiosis to koinobiont endoparasitism, as suggested by (Gauld, 1988), or it could represent a reversal in this lineage to ectoparasitism (Quicke, 2015).

The results presented here suggests that strong conclusions cannot be drawn from these data. Teasing apart the full nature of how different parasitoid strategies evolved will require a very detailed look at host taxa and host biology and the stage attacked (Quicke et al., 1999a; Quicke, 2015), which is sorely lacking for most Ichneumonoidea. Although the sampling is likely too low to draw strong conclusions about the number of transitions across Braconidae and Ichneumonidae, there certainly have been transitions from koinobiosis back to idiobiosis and from ecto– to endoparasitism in both families. This was advocated as well by Bennett et al. (2019) who also suggested mechanisms for transition from koinobiosis back to idiobiosis, challenging the prevailing notion that this transition was rare or non-existent (Gauld, 1988; Quicke, 2015; Quicke et al., 2000).

### 4.4 Evolution of Endogenous Virus Elements in Wasp Genomes

Endogenous virus elements that produce virions (PDVs) or VLPs have been described in both braconids and ichneumonids. Until recently, Braconidae was thought to have a single origin of endogenous viruses in the microgastroid lineage (Beckage et al., 1994; Bézier et al., 2009; Huguet et al., 2012; Strand and Burke, 2015; Whitfield, 1997, 2002). A new endogenous virus descended from pathogenic alphanudiviruses was discovered in some species of *Fopius*, opiine braconids, representing a second integration event and a relatively recent one (Burke, 2019; Burke et al., 2018a). In Ichneumonidae, PDVs with separate evolutionary origins from braconid PDVs have been discovered in members of Banchinae and Campopleginae (Espagne et al., 2004; Lapointe et al., 2007; Norton et al., 1975; Robin et al., 2019; Stoltz et al., 1981; Stoltz and Whitfield, 1992; Tanaka et al., 2007; Volkoff et al., 2010). Further, at least one species of campoplegine has acquired a VLP related to nudiviruses (Pichon et al., 2015; Reineke et al., 2006; Rotheram, 1973). Given the robust phylogeny here, it is clear that banchines and campoplegines are not closely related, and the intervening taxa have not been verified to possess PDVs (but see Cummins et al., 2011). Although the ichnoviruses in Campopleginae and Banchinae come from closely related virus ancestors (Béliveau et al., 2015), our data here provide strong evidence that the Campopleginae and Banchinae are not sister subfamilies within the Ichneumonidae. Two possibilities exist: first, that there have been two independent origins of ichnoviruses in the Ichneumonidae, or second, that there was a single origin and ichnoviruses were lost in some other subfamilies between the ichnovirus-producing families (Beliveau et al. 2015). In a companion study (Burke et al., 2020), a more extensive search for PDVs in intervening taxa using a genome sequencing approach revealed that there are no ichnovirus genes in several intervening taxa, consistent with the failure to capture ichnovirus genes within these taxa using the anchored hybrid enrichment approach used in this study.

PDV genes are known to evolve very rapidly, presenting challenges for finding these genes in the genome with targeted approaches such as PCR. Targeted enrichment approaches hold some promise for sampling across a broader array of PDV genes specific to certain taxa to determine whether they may be present in previously untested species and how these genes have evolved. We have found ichnovirus genes in the *Hyposoter* and *Campoletis* sampled here, but they are not identified to species. However, *Campoletis* sp. 1 was collected in Australia, where *C. sonorensis* is not known to occur, and thus represents at least one new species with an ichnovirus in this genus. Further, we provide the first evidence of PDVs in the genus *Casinaria*. The gene used, IVSP4, appears to undergo within lineage gene expansions or contractions, which the AHE pipelines were able to detect, but will require more data from more taxa to better understand their evolution. These genes were not captured in species of *Dusona* and *Campoplex*, which were recovered as sister taxa, which suggests these taxa either do not have ichnoviruses or they have very divergent copies. Without richer taxonomic sampling their sister relationship cannot be certain. In another recent study, *Dusona* was recovered as sister to *Casinaria* (Bennett et al., 2019), and a study with deeper sampling recovered conflicting placement and non-monophyly for several campoplegine genera (Quicke et al., 2009).

For the bracovirus replication gene *HzNVorf128* we were able to fully capture the gene from all expected bracovirus-harboring taxa and reconstruct a pattern of evolution that is consistent with species relationships determined using other non-viral molecular markers. Thus, capture in *Phanerotoma* and *Schoenlandella* represent confirmations in new taxa, but these were expected given that microgastroids are all assumed to have bracoviruses (Whitfield, 1997, 2002). However, the other bracovirus gene (*per os infectivity factor p74)* was not captured in full across likely bracovirus harboring taxa, most likely due to a more rapid evolutionary rate. In general, these genes appear to be evolving rapidly, thus limiting the ability to capture some viral genes using probes from known sequences. It is conceivable that more targeted approaches that iteratively design probes focused upon single-copy genes with high sequence conservation may be more effective for cross-lineage capture of divergent viral genes.

## 5. Conclusions

Here we developed a 541 gene probe set for anchored hybrid enrichment that was largely successful for examining relationships in Ichneumonoidea and with wide utility across Hymenoptera. Although we also targeted polydnavirus genes (PDVs), capture was not entirely successful across taxa known to have viruses endogenized in their genomes. It is possible that a more lineage-specific targeted approach to probe design could be more successful, but genome sequencing may be a more appropriate method for discovering virus association and endogenization given the continually decreasing cost. Higher-level relationships across Ichneumonoidea are mostly stable and well supported relationships and reinforce various findings from other studies. While we focused on Ichneumonidae, the braconid relationships were largely consistent with previous studies. Within Ichneumonidae, Xoridinae was the most basal taxon, and the placements of Labeninae and Orthopelmatinae remain uncertain. Excluding these three taxa, we recovered Ophioniformes (Ichneumoniformes (including Eucerotinae) + Pimpliformes). Higher-level relationships among Ichneumonoidea have been historically difficult to resolve, but anchored hybrid enrichment offers a valuable tool to recovering robust relationships in this highly diverse lineage. However, our data demonstrate that systematic biases need to be thoroughly investigated to prevent erroneous relationships and outgroup selection must be vetted. We found codon use bias present in some lineages, suggesting amino acid data may be better suited to understanding evolution in Ichneumonoidea.

Ancestral state reconstructions were sensitive to the taxa included or data used to infer the character states. The data suggest that the ancestor of Ichneumonoidea was an idiobiont ectoparasitoid. While analyses for Ichneumonidae were stable for idiobiosis/koinobiosis, they varied for ecto- versus endoparasitism. Braconidae may have had a koinobiont endoparasitoid ancestor, although reconstructions varied widely for the family preventing strong conclusions. The results suggest many transitions of these traits within Ichneumonoidea and full understanding of the evolution of parasitoid strategies will require deep taxonomic sampling and increased knowledge on the biology of this diverse lineage. At least two independent acquisitions of endogenous polydnaviruses (PDVs) within Ichneumonidae are well supported by the data due to the distant relationship between Banchinae and Campopleginae and lack of evidence of PDVs in intervening taxa. However, probes designed to capture polydnavirus genes likely require a more targeted lineage-specific approach.

## Acknowledgements

We are deeply thankful for specimens from Michael Sharkey, David Smith, and Dave Karlsson with the Swedish Malaise Trap Project (SMTP). We thank Geoff Allen (University of Tasmania), Andrew Austin (University of Adelaide), James Adams (La Ceiba, Honduras), and Bery Josue Almendares Romero and entomological researchers at the Universidad Nacional Autónoma de Honduras for hosting during collecting trips and assisting with permits. Our grateful thanks to Davide Dal Pos and Andrés Herrera-Flórez for some specimen identifications and to Davide Dal Pos for valuable comments on manuscript drafts. A special thank-you to Andrew Bennett for many discussions about relationships across the Ichneumonidae. We thank Terry Wheeler (deceased) and Chris Buddle for organizing a collecting trip to Yukon through the Northern Biodiversity Program (Canada). We would like to thank John Heraty for providing alignments for the probe design, which increased their utility across Hymenoptera. We are grateful to Michelle Kortyna and Alyssa Bigelow at FSU’s Center for Anchored Phylogenomics for assistance with sample processing. Funding support for this project was provided by the Natural Sciences and Engineering Research Council (NSERC) Discovery Program to BJS and the National Science Foundation (NSF) (Award Number: 1916914) to BJS and GRB.

