## Supplemental Figures and Tables for "Phylogenomics of Ichneumonoidea (Hymenoptera) and implications for evolution of mode of parasitism and viral endogenization"

### Table of Contents

**SI Tables**

**Table S1. Character coding for ancestral state reconstructions** ……………………. 4

**Table S2**. **Taxon exclusion tests** ………………………………………………………… 6

**SI Figures**

**Figure S1. Summary of sequence assembly results** ………………….……………….. 7

**Figure S2**. **Saturation plots by codon position** …………………………………………. 8

**Figure S3. Maximum-likelihood (ML) phylogeny of Ichneumonoidea based on 479 genes analyzed as nucleotides** ………………………………………………………………… 9

**Figure S4**. **Maximum-likelihood (ML) phylogeny of Ichneumonoidea based on 479 genes analyzed as nucleotides with the 3rd position removed (nt-3out)** ……….…………. 10

**Figure S5. Bayesian phylogeny of Ichneumonoidea based on 479 genes analyzed as nucleotides with the 3rd position removed** ……………..…………………………….. 11

**Figure S6. Maximum-likelihood (ML) phylogeny of Ichneumonoidea based on 479 genes analyzed as amino acids with outgroups excluded** ……………………..................... 12

**Figure S7. Maximum-likelihood (ML) phylogeny of Ichneumonoidea based on 479 genes analyzed as amino acids with Aphidiinae excluded** …………………………………… 13

**Figure S8. Maximum-likelihood (ML) phylogeny of Ichneumonoidea based on 479 genes analyzed as amino acids with Agathidinae excluded** ………………………………..... 14

**Figure S9. Maximum-likelihood (ML) phylogeny of Ichneumonoidea based on 479 genes analyzed as amino acids with Xoridinae excluded** ……………………………….......... 15

**Figure S10. Maximum-likelihood (ML) phylogeny of Ichneumonoidea based on 479 genes analyzed as amino acids with Labeninae excluded** ……………….......................... 16

**Figure S11. Maximum-likelihood (ML) phylogeny of Ichneumonoidea based on 479 genes analyzed as amino acids with Orthopelmatinae excluded** …………………………....... 17

**Figure S12. Maximum-likelihood (ML) phylogeny of Ichneumonoidea based on 479 genes analyzed as amino acids with Labeninae and Orthopelmatinae excluded** ………... 18

**Figure S13. Maximum-likelihood (ML) phylogeny of Ichneumonoidea based on 479 genes analyzed as amino acids with Eucerotinae excluded** …………………...................... 19

**Figure S14. Maximum-likelihood (ML) phylogeny of Ichneumonoidea based on 479 genes analyzed as amino acids with Adelognathinae excluded** ………………….................. 20

**Figure S15. Maximum-likelihood (ML) phylogeny of Ichneumonoidea based on 479 genes analyzed as amino acids with Hybrizontinae excluded** ……………………............. 21

**Figure S16. Maximum-likelihood (ML) phylogeny of Ichneumonoidea based on 479 genes analyzed as amino acids with Mesochorinae excluded** …………….......................... 22

**Figure S17. Maximum-likelihood (ML) phylogeny of Ichneumonoidea based on 479 genes analyzed as amino acids with Tersilochinae excluded** …………………………......... 23

**Figure S18. Principle Component Analysis (PCA) of Relative Synonymous Codon Usage (RSCU) between select major groups** ……………………..………………………………… 24

**Figure S19. Principle Component Analysis (PCA) of Relative Synonymous Codon Usage (RSCU) between select major groups and sampled Pimplinae tribes (ePhialtini and Pimplini)** …………………………………………………………………………………...…. 25

**Table S1**: **Character coding for ancestral state reconstructions.**

| **Taxon** | **Character 1**: Idiobiont (0); Koinobiont (1) | **Character 2**: Ectoparasitoid (0); Endoparasitoid (1) |
|---|---|---|
| I10000_Ctenopelmatinae_Megaceria_sp | 1 | 1 |
| I10005_Campopleginae_Casinaria_sp1 | 1 | 1 |
| I10007_Campopleginae_Campoletis_sp1 | 1 | 1 |
| I2339_Campopleginae_Campoletis_sp2 | 1 | 1 |
| I9995_Campopleginae_Hyposoter_sp* | 1 | 1 |
| I10006_Campopleginae_Campoplex_sp1* | 1 | 1 |
| I9991_Campopleginae_Campoplex_sp2 | 1 | 1 |
| I10011_Campopleginae_Dusona_sp3* | 1 | 1 |
| I10012_Campopleginae_Dusona_sp1 | 1 | 1 |
| I15703_Campopleginae_Dusona_sp2 | 1 | 1 |
| I2327_Ophioninae_Ophion_sp | 1 | 1 |
| I10009_Anomaloninae_Habronyx_sp | 1 | 1 |
| I2337_Anomaloninae_Agrypon_sp | 1 | 1 |
| I2336_Cremastinae_Cremastus_sp | 1 | 1 |
| I9981_Cremastinae_Pristomerus_sp | 1 | 1 |
| I9982_Hybrizontinae_Hybrizon_buccatus | 1 | 1 |
| I2334_Metopiinae_Exochus_sp | 1 | 1 |
| I2357_Metopiinae_Spudaeus_sp | 1 | 1 |
| I9993_Metopiinae_Metopius_sp | 1 | 1 |
| I15696_Ctenopelmatinae_Anoncus_sp | 1 | 1 |
| I2340_Ctenopelmatinae_Mesoleius_sp1* | 1 | 1 |
| I9985_Oxytorinae_Oxytorus_sp | ? | ? |
| I15709_Mesochorinae_Mesochorus_sp | 1 | 1 |
| I2341_Tersilochinae_Phradis_sp1* | 1 | 1 |
| I9992_Tersilochinae_Stethantyx_sp | 1 | 1 |
| I10001_Banchinae_Meniscomorpha_sp1 | 1 | 1 |
| I10003_Banchinae_Meniscomorpha_sp2 | 1 | 1 |
| I15708_Banchinae_Lissonota_sp | 1 | 1 |
| I10008_Banchinae_Australoglypta_sp | 1 | 1 |
| I10013_Banchinae_Lissonota_sp1* | 1 | 1 |
| I2338_Banchinae_Glypta_sp | 1 | 1 |
| I2324_Tryphoninae_Exenterus_sp | 1 | 0 |
| I2335_Tryphoninae_Netelia_Netelia_sp | 1 | 0 |
| I9983_Tryphoninae_Phytodietus_sp | 1 | 0 |
| I10004_Pimplinae_Eriostethus_pulcherrimus | 1 | 0 |
| I2356_Pimplinae_Scambus_sp | 0 | 0 |
| I15702_Pimplinae_Dolichomitus_sp | 0 | 0 |
| I2330_Pimplinae_Pimpla_pedalis | 0 | 1 |
| I9996_Pimplinae_Pimpla_molesta | 0 | 1 |
| I9998_Pimplinae_Echthromorpha_intricatoria | 0 | 1 |
| I2331_Poemeniinae_Neoxorides_borealis | 0 | 1 |
| I2332_Diplazontinae_Homotropus_sp | 1 | 1 |
| I2333_Orthocentrinae_Orthocentrus_sp | 1 | 1 |

| | | |
|---|---|---|
| I9987_Collyriinae_Collyria_sp | 1 | 1 |
| I9988_Cylloceriinae_Cylloceria_sp | ? | 1 |
| I2326_Acaenitinae_Spilopteron_occiputale | 1 | 1 |
| I10010_Ichneumoninae_Ichneumonini | 1 | 1 |
| I9989_Ichneumoninae_Akymichneumon_sp | 1 | 1 |
| I9999_Ichneumoninae_Gavrana_maculipes_sg | 1 | 1 |
| I9997_Ichneumoninae_Gavrana_sp | 1 | 1 |
| I2328_Cryptinae_Mesostenus_thoracicus | 0 | 1 |
| I2329_Cryptinae_Echthrus_abdominalis | 0 | 0 |
| I15694_Adelognathinae_Adelognathus_sp | 0 | 0 |
| I9986_Eucerotinae_Euceros_sp | 1 | 1 |
| I9984_Orthopelmatinae_Orthopelma_sp | 1 | 1 |
| I10002_Labeninae_Labium_sp | 0 | 0 |
| I9990_Labeninae_Certonotus_nitidulus1 | 0 | 0 |
| I9994_Labeninae_Certonotus_nitidulus2 | 0 | 0 |
| I15713_Xoridinae_Odontocolon_sp | 0 | 0 |
| I15695_Rogadinae_Aleiodes_sp | 1 | 1 |
| I15701_Opiinae_Diachasma_alloeum | 1 | 1 |
| I2359_Alysiinae_Chorebus_sp | 1 | 1 |
| I2355_Braconinae_Bracon_sp | 0 | 0 |
| I2358_Pambolinae_Pambolus_sp | 0 | 0 |
| I2325_Rhyssalinae_Rhyssalus_sp | 0 | 0 |
| I2347_Aphidiinae_Pauesia_sp | 1 | 1 |
| I2352_Aphidiinae_Praon_sp | 1 | 1 |
| I15704_Helconinae_Eumacrocentrus_americanus | 1 | 1 |
| I2342_Orgilinae_Orgilus_ablusus* | 1 | 1 |
| I2345_Brachistinae_Blacus_sp | 1 | 1 |
| I2353_Brachistinae_Diospilus_sp | 1 | 1 |
| I2346_Brachistinae_Apoblacus_sp | 1 | 1 |
| I2354_Acampsohelconinae_Urosigalphus_sp | 1 | 1 |
| I15710_Euphorinae_Meterous_sp | 1 | 1 |
| I15711_Microgastrinae_Microplitis_demolitor | 1 | 1 |
| I2349_Cardiochilinae_Schoenlandella_sp | 1 | 1 |
| I15714_Cheloninae_Phanerotoma_sp | 1 | 1 |
| I2351_Cheloninae_Chelonus_sp | 1 | 1 |
| I2344_Agathidinae_Bassus_sp | 1 | 1 |
| I2343_Ichneutinae_Ichneutes_sp | 1 | 1 |
| I15693_Formicidae_Acromyrmex_echinatior | - | - |
| I15698_Formicidae_Atta_cephalotes | - | - |
| I15716_Formicidae_Solenopsis_invicta | - | - |
| I15715_Formicidae_Pogonomyrmex_barbatus | - | - |
| I15700_Formicidae_Camponotus_floridanus | - | - |
| I15706_Formicidae_Linepithema_humile | - | - |
| I15705_Formicidae_Harpegnathos_saltator | - | - |
| I15697_Apidae_Apis_mellifera | - | - |
| I15699_Apidae_Bombus_impatiens | - | - |
| I15707_Megachilidae_Megachile_rotunda | - | - |
| I15712_Pteromalidae_Nasonia_vitripennis | 0 | 1 |

**Table S2**: **Taxon exclusion tests**. Summary of select clades recovered after phylogenetic analysis with select taxa excluded. Listed supplementary figures depict the entire phylogeny for the specific taxa removed. Increases and decreases in support values (into new categories, as per legend in Fig. 1a-c) are indicated by a "+" or "-," respectively; LRT values shown in first cell and UFb values shown in second cell. Xor =Xoridinae; Try = Tryphoninae.

| Clade (Fig. 1a-c) | **Supplementary Figure** / **Description of Clade** | Fig. S6 | | Fig. S7 | | Fig. S8 | | Fig. S9 | |
|---|---|---|---|---|---|---|---|---|---|
| | | OGs removed; OG = Rhyssalinae; subset of Braconids | | Aphidiinae removed | | Agathidinae removed | | Xoridinae removed | |
| A | Rogadinae sister to Alysioid subcomplex | **Y+** | **Y** | **Y+** | **Y** | **Y** | **Y** | **Y** | **Y** |
| B | Agathidinae sister to Microgastroid complex | **N/A** | **N/A** | **Y** | **Y** | **N/A[a]** | **N/A[a]** | **Y** | **Y** |
| C | Labeninae sister to all Ichneumonidae (except Xor) | **N** | **N** | **Y** | **Y** | **Y** | **Y** | **Y+** | **Y** |
| D | Orthopelmatinae sister to Ichneumoniformes *s.l.* | **Y+** | **Y** | **Y** | **Y** | **Y** | **Y** | **Y** | **Y** |
| E | Banchinae sister to all Ophioniformes (except Try) | **Y+** | **Y** | **Y** | **Y** | **Y** | **Y** | **Y** | **Y** |
| | Pimpliformes relationships | **same** | | **same** | | **same** | | **same** | |

| Clade (Fig. 1a-c) | **Supplementary Figure** / **Description of Clade** | Fig. S10 | | Fig. S11 | | Fig. S12 | | Fig. S13 | |
|---|---|---|---|---|---|---|---|---|---|
| | | Labeninae removed | | Orthopelmatinae removed | | Labeninae + Orthopelmatinae removed | | Eucerotinae removed | |
| **A** | Rogadinae sister to Alysioid subcomplex | **Y** | **Y** | **Y** | **Y** | **Y** | **Y** | **Y** | **Y** |
| B | Agathidinae sister to Microgastroid complex | **Y** | **Y** | **Y** | **Y** | **Y** | **Y** | **Y** | **Y** |
| C | Labeninae sister to all Ichneumonidae (except Xor) | **N/A[b+]** | **N/A[b]+** | **Y** | **Y** | **N/A[d]** | **N/A[d]** | **Y** | **Y-** |
| D | Orthopelmatinae sister to Ichneumoniformes *s.l.* | **Y** | **Y-** | **N/A** | **N/A** | **N/A[d]** | **N/A[d]** | **Y+** | **Y+** |
| E | Banchinae sister to all Ophioniformes (except Try) | **Y** | **Y** | **Y** | **Y** | **Y** | **Y** | **Y** | **Y** |
| | Pimpliformes relationships | **same** | | **same[c]** | | **same** | | **same** | |

| Clade (Fig. 1a-c) | **Supplementary Figure** / **Description of Clade** | Fig. S14 | | Fig. S15 | | Fig. S16 | | Fig. S17 | |
|---|---|---|---|---|---|---|---|---|---|
| | | Adelognathinae removed | | Hybrizontinae removed[f] | | Mesochorinae removed | | Tersilochinae removed | |
| **A** | Rogadinae sister to Alysioid subcomplex | **Y** | **Y** | **Y** | **Y** | **Y** | **Y** | **Y** | **Y** |
| B | Agathidinae sister to Microgastroid complex | **Y** | **Y** | **Y** | **Y** | **Y** | **Y** | **Y** | **Y** |
| C | Labeninae sister to all Ichneumonidae (except Xor) | **N[e]** | **N[e]** | **Y** | **Y** | **Y** | **Y** | **N[h]** | **N[h]** |
| D | Orthopelmatinae sister to Ichneumoniformes *s.l.* | **Y+** | **Y-** | **Y** | **Y** | **Y** | **Y** | **Y+** | **Y** |
| E | Banchinae sister to all Ophioniformes (except Try) | **Y** | **Y** | **Y** | **Y** | **N[g]** | **N[g]** | **N/A** | **N/A** |
| | Pimpliformes relationships | **same** | | **same** | | **same** | | **same** | |

[a]Support for Ichneutinae + Microgastroid complex increases when Agathidinae is removed.

[b]Clade containing all Ichneumonidae except Xoridinae and Labeninae (although excluded) increases substantially.

[c]When Orthopelmatinae is removed, the relationship of Diplazontinae + Orthocentrinae as sister to Poeminae + Pimplinae is substantially decreased

[d]When both these taxa are removed, the relationship of Ophioniformes (Ichneumoniformes + Pimpliformes) becomes >99 for all clades.

[e]Labeninae is recovered as sister to Pimpliformes with very low support (< 35).

[f]Anomalinae is recovered as sister to Cremastinae with relatively high support (>93)

[g]Banchinae is recovered as sister to Tersilochinae, similar to the nt-3out analyses (Figs. S4-5), albeit with low support (12/63)

[h]Labeninae is recovered as sister to Orthopelmatinae + Ichneumoniformes *s.l.* with very low support (6/39)

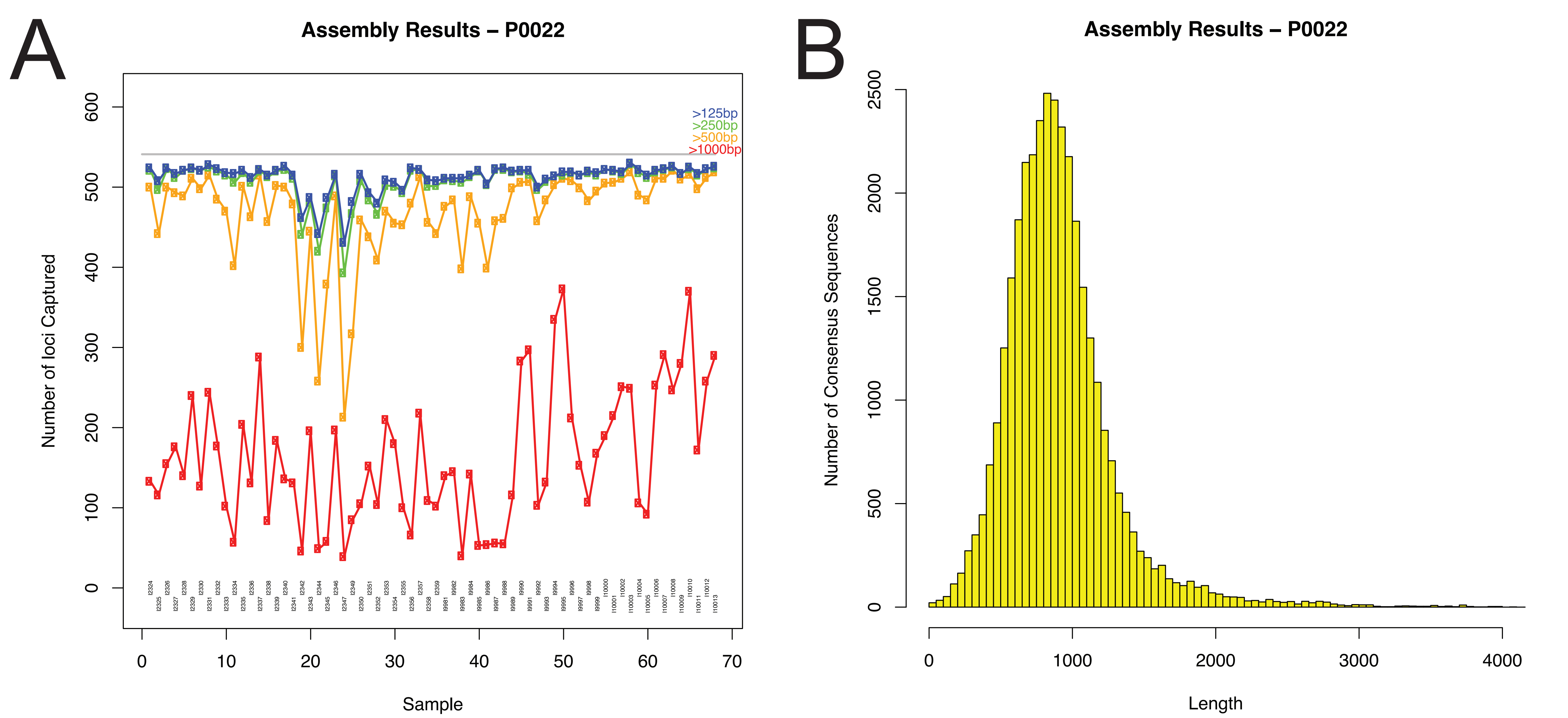


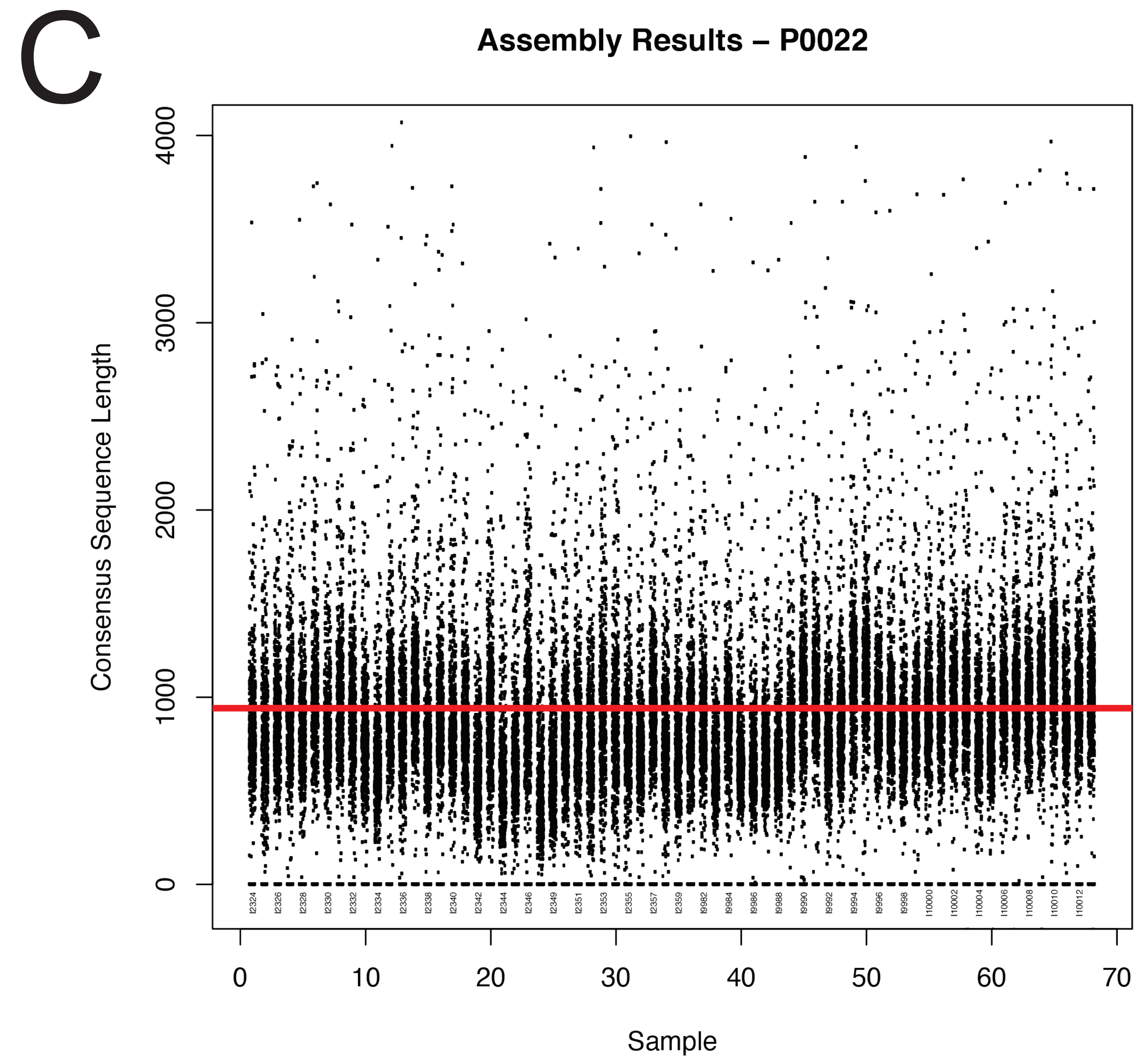


**Figure S1. Summary of sequence assembly results**. A. Number of loci captured per consensus sequence size. B. Histogram of sequence length for consensus sequences. C. Dotplot of consensus sequence length by taxon, red line indicates average length.

a. First Codon Position

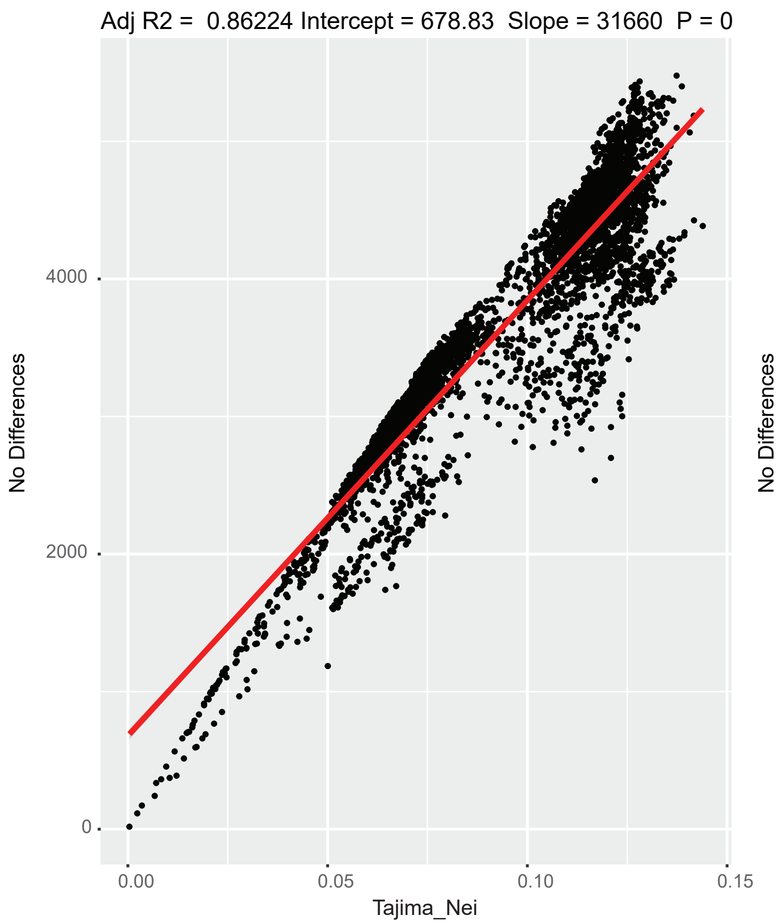


b. Second Codon Position

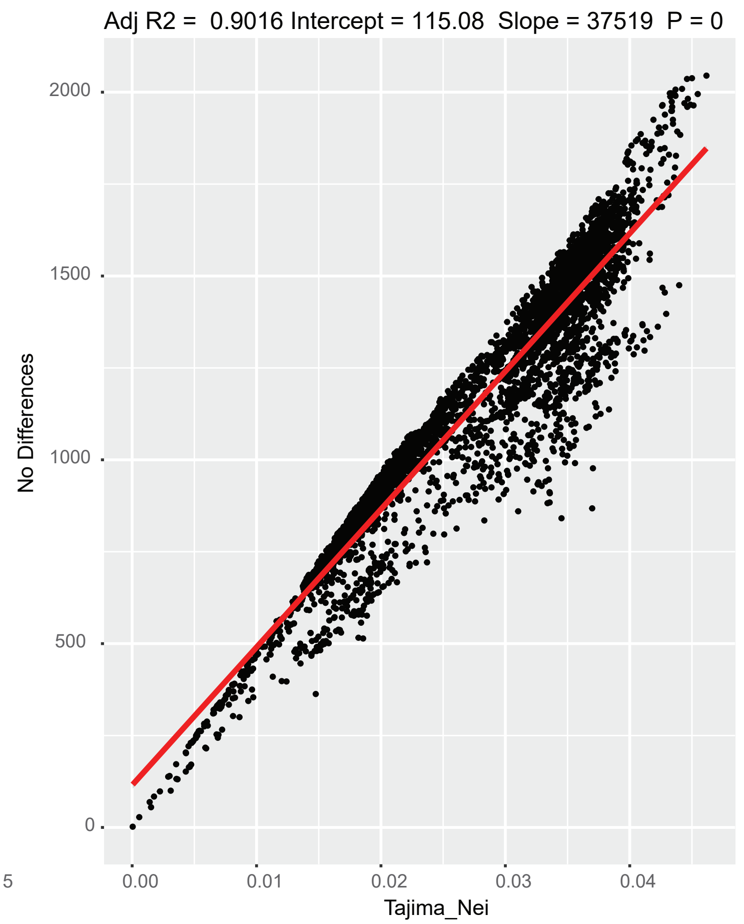


c. Third Codon Position

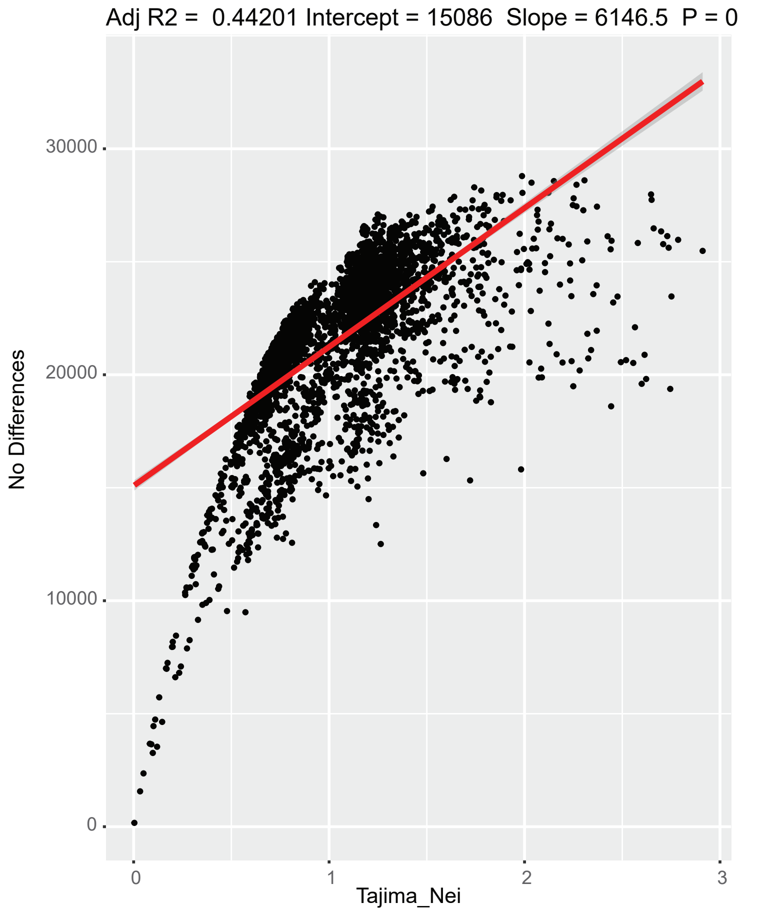


**Figure S2. Saturation plots by codon position.** Tajima-Nei genetic distance is plotted against the total number of differences in pairwise alignments. a. First codon position; b. Second codon position; c. Third codon position.

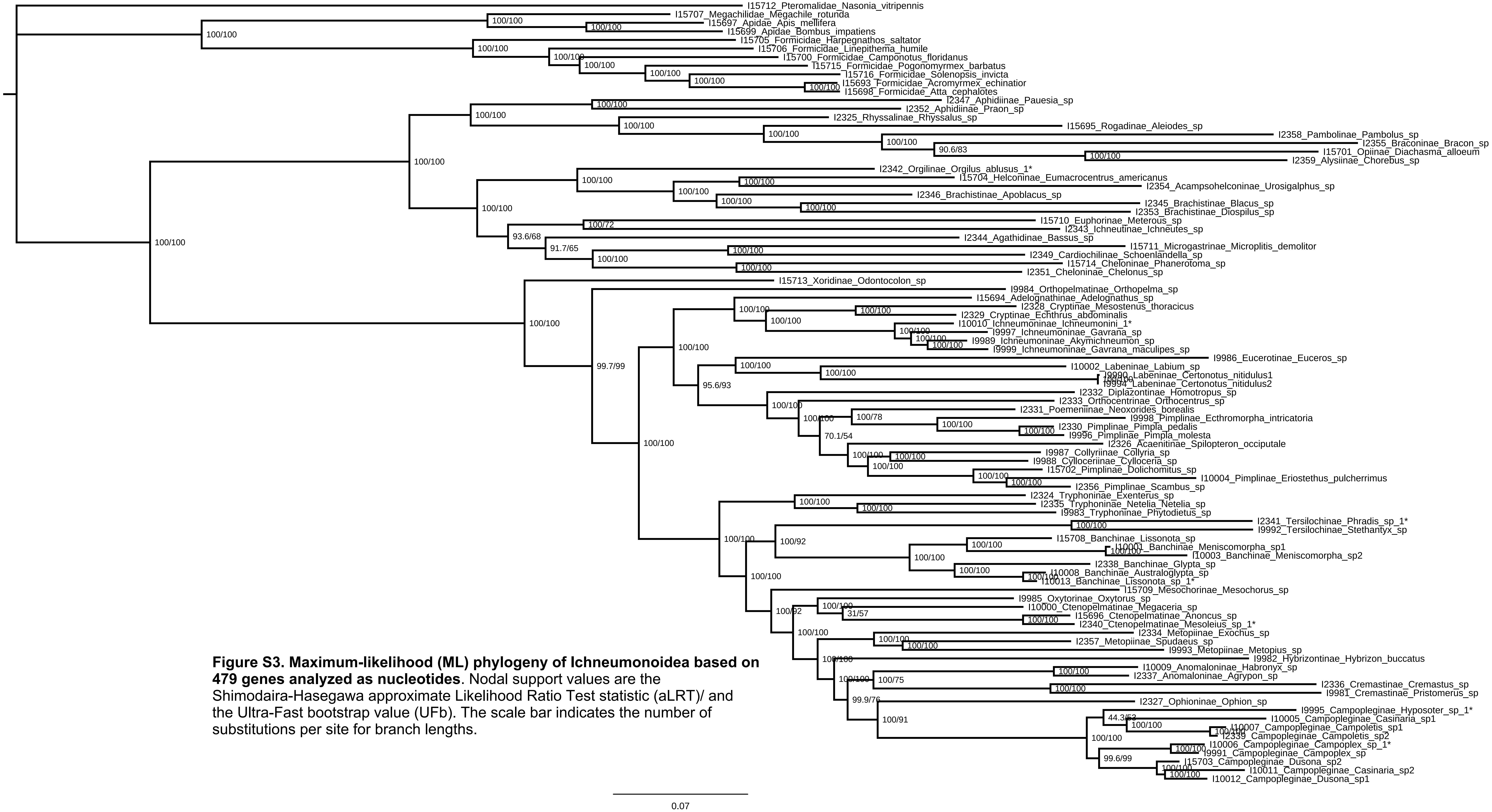


**Figure S3. Maximum-likelihood (ML) phylogeny of Ichneumonoidea based on 479 genes analyzed as nucleotides**. Nodal support values are the Shimodaira-Hasegawa approximate Likelihood Ratio Test statistic (aLRT)/ and the Ultra-Fast bootstrap value (UFb). The scale bar indicates the number of substitutions per site for branch lengths.

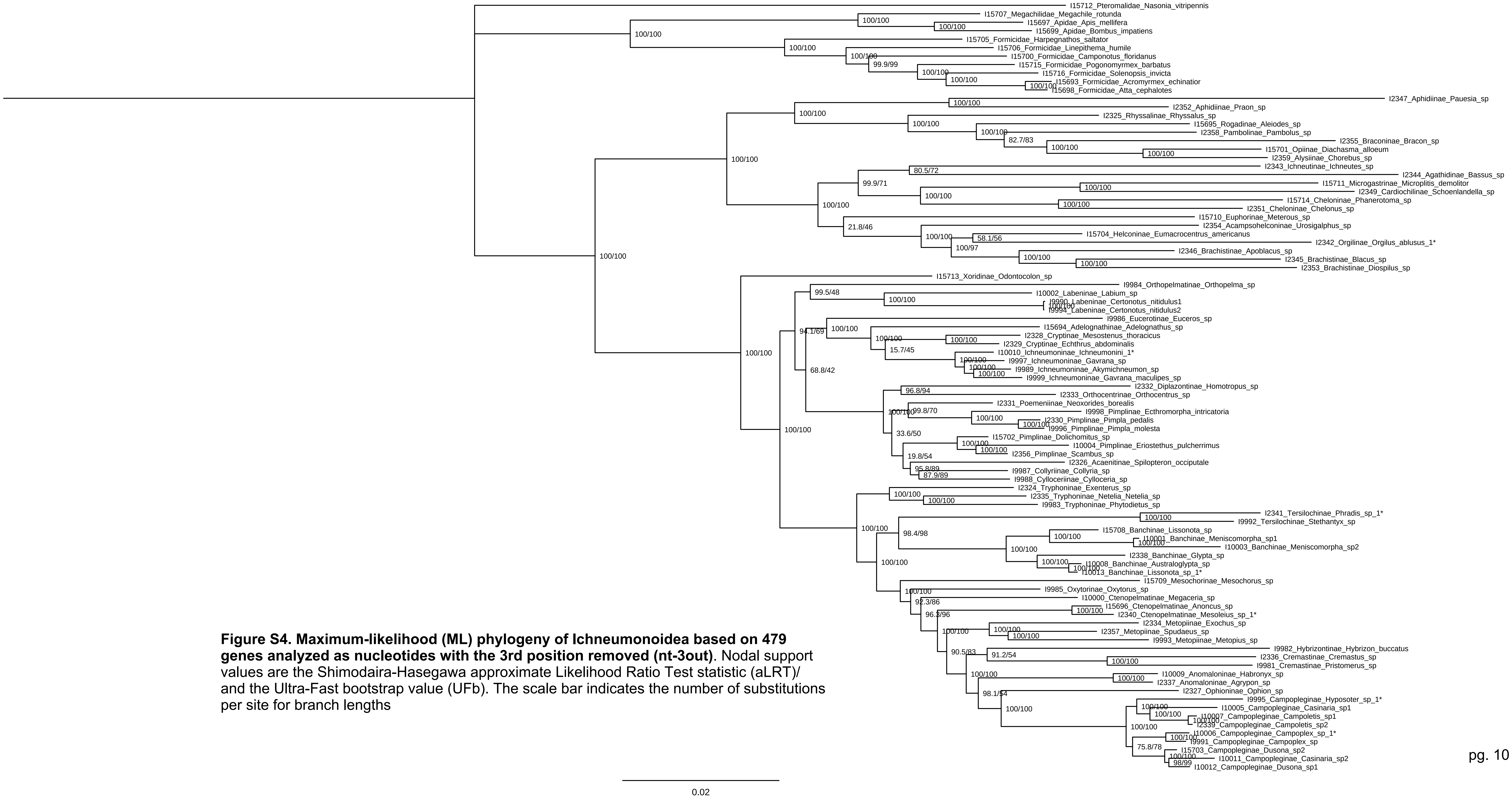


**Figure S4. Maximum-likelihood (ML) phylogeny of Ichneumonoidea based on 479 genes analyzed as nucleotides with the 3rd position removed (nt-3out)**. Nodal support values are the Shimodaira-Hasegawa approximate Likelihood Ratio Test statistic (aLRT)/ and the Ultra-Fast bootstrap value (UFb). The scale bar indicates the number of substitutions per site for branch lengths

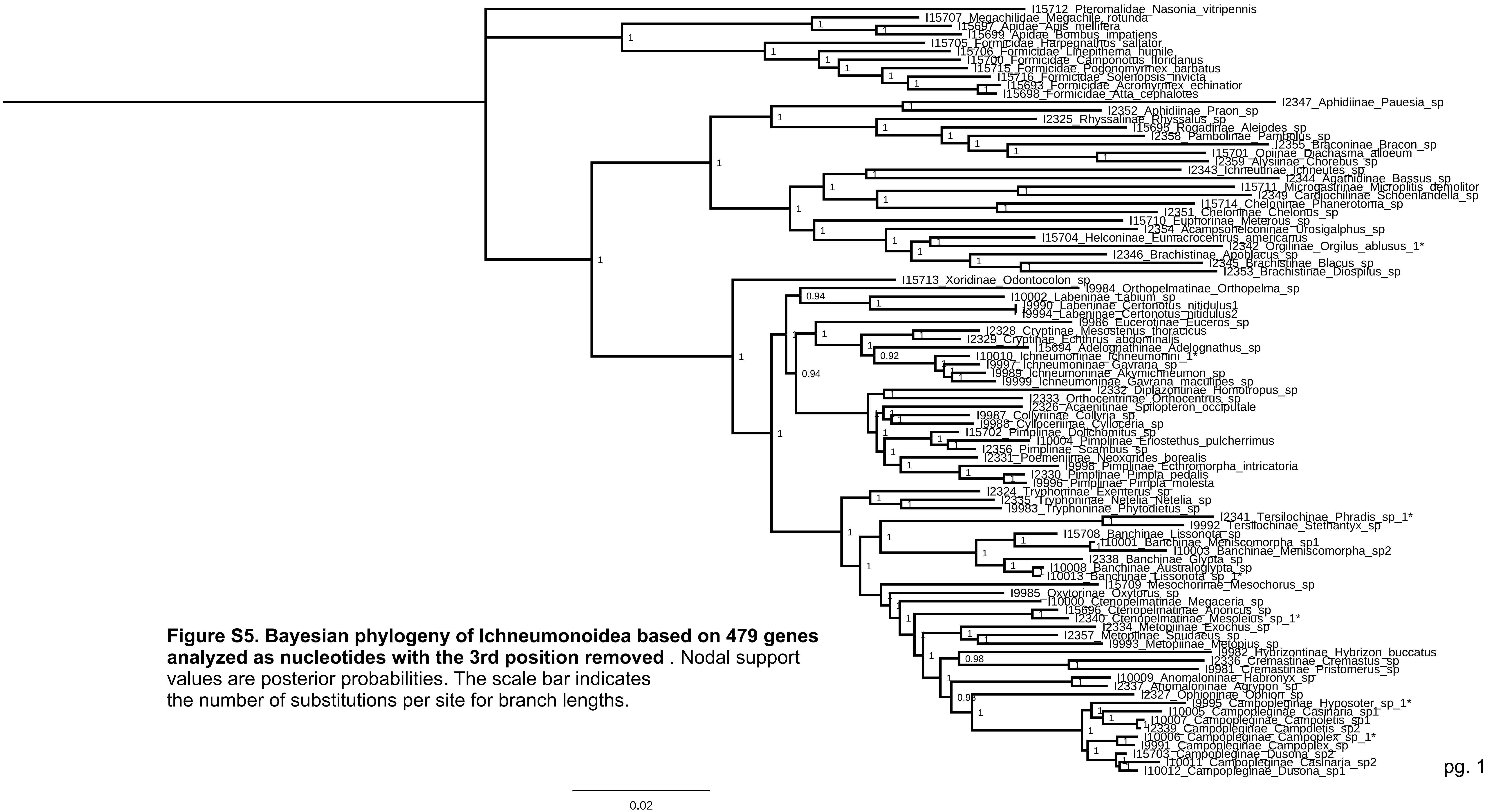


**Figure S5. Bayesian phylogeny of Ichneumonoidea based on 479 genes analyzed as nucleotides with the 3rd position removed** . Nodal support values are posterior probabilities. The scale bar indicates the number of substitutions per site for branch lengths.

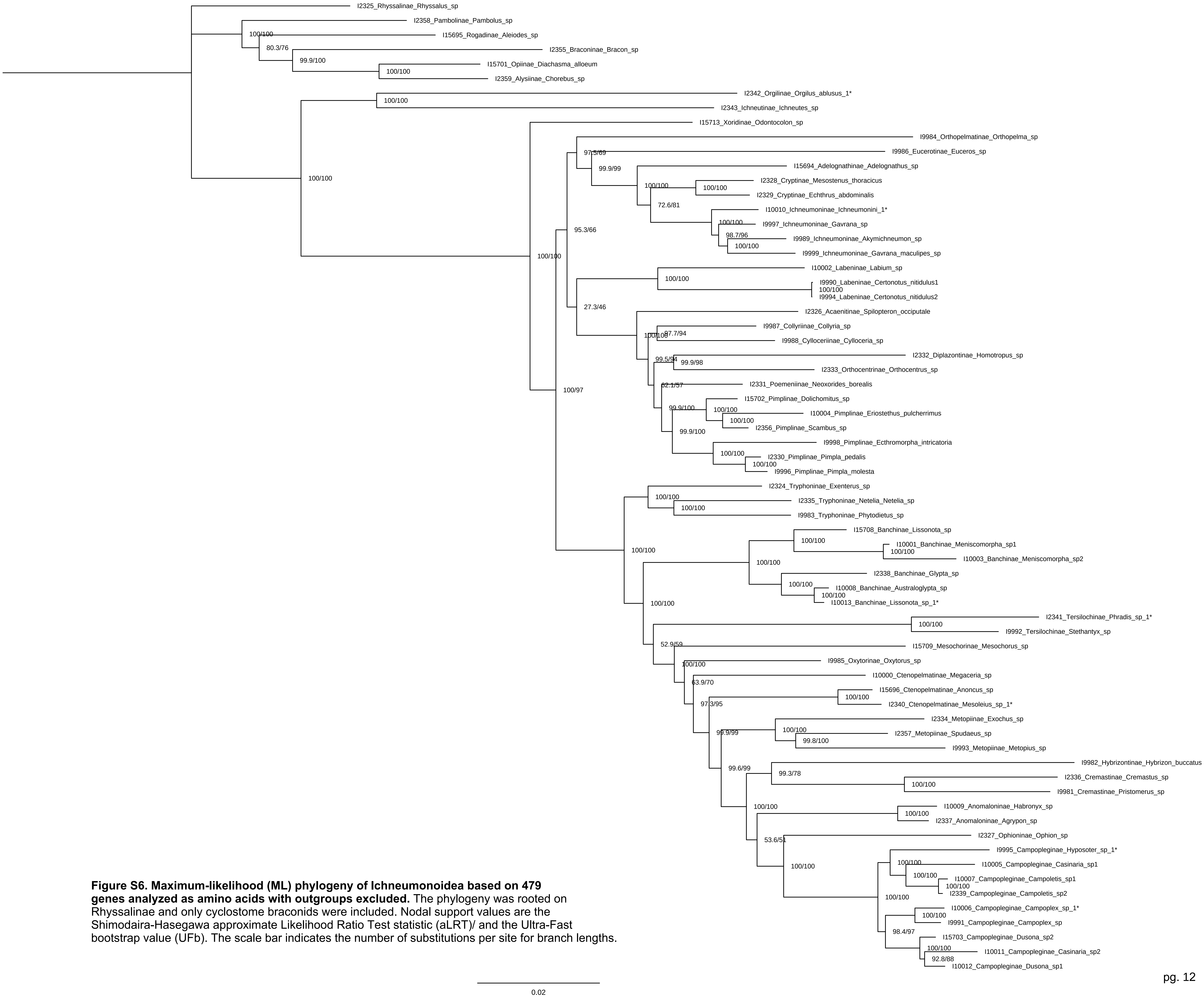


**Figure S6. Maximum-likelihood (ML) phylogeny of Ichneumonoidea based on 479 genes analyzed as amino acids with outgroups excluded.** The phylogeny was rooted on Rhyssalinae and only cyclostome braconids were included. Nodal support values are the Shimodaira-Hasegawa approximate Likelihood Ratio Test statistic (aLRT)/ and the Ultra-Fast bootstrap value (UFb). The scale bar indicates the number of substitutions per site for branch lengths.

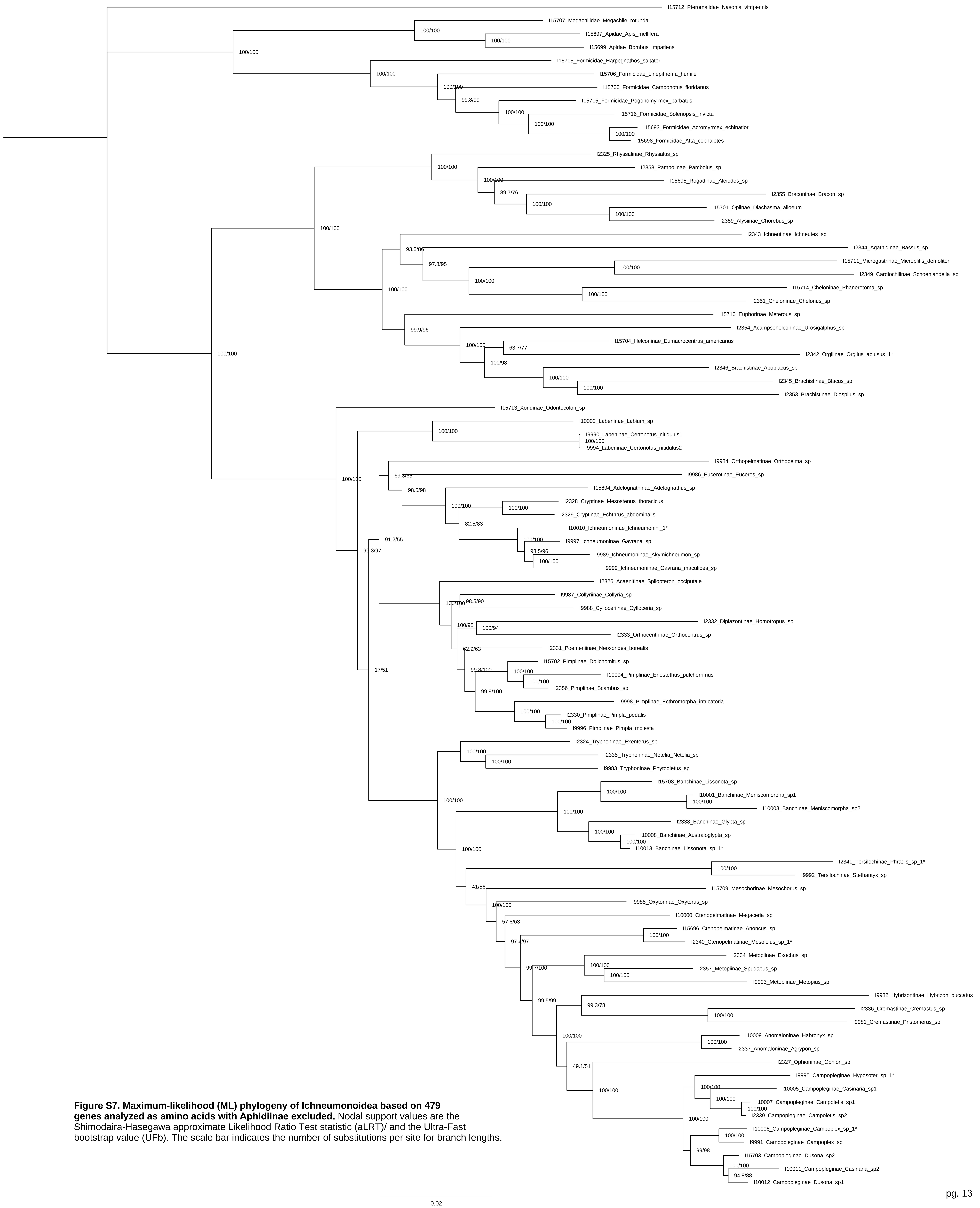


**Figure S7. Maximum-likelihood (ML) phylogeny of Ichneumonoidea based on 479 genes analyzed as amino acids with Aphidiinae excluded.** Nodal support values are the Shimodaira-Hasegawa approximate Likelihood Ratio Test statistic (aLRT)/ and the Ultra-Fast bootstrap value (UFb). The scale bar indicates the number of substitutions per site for branch lengths.

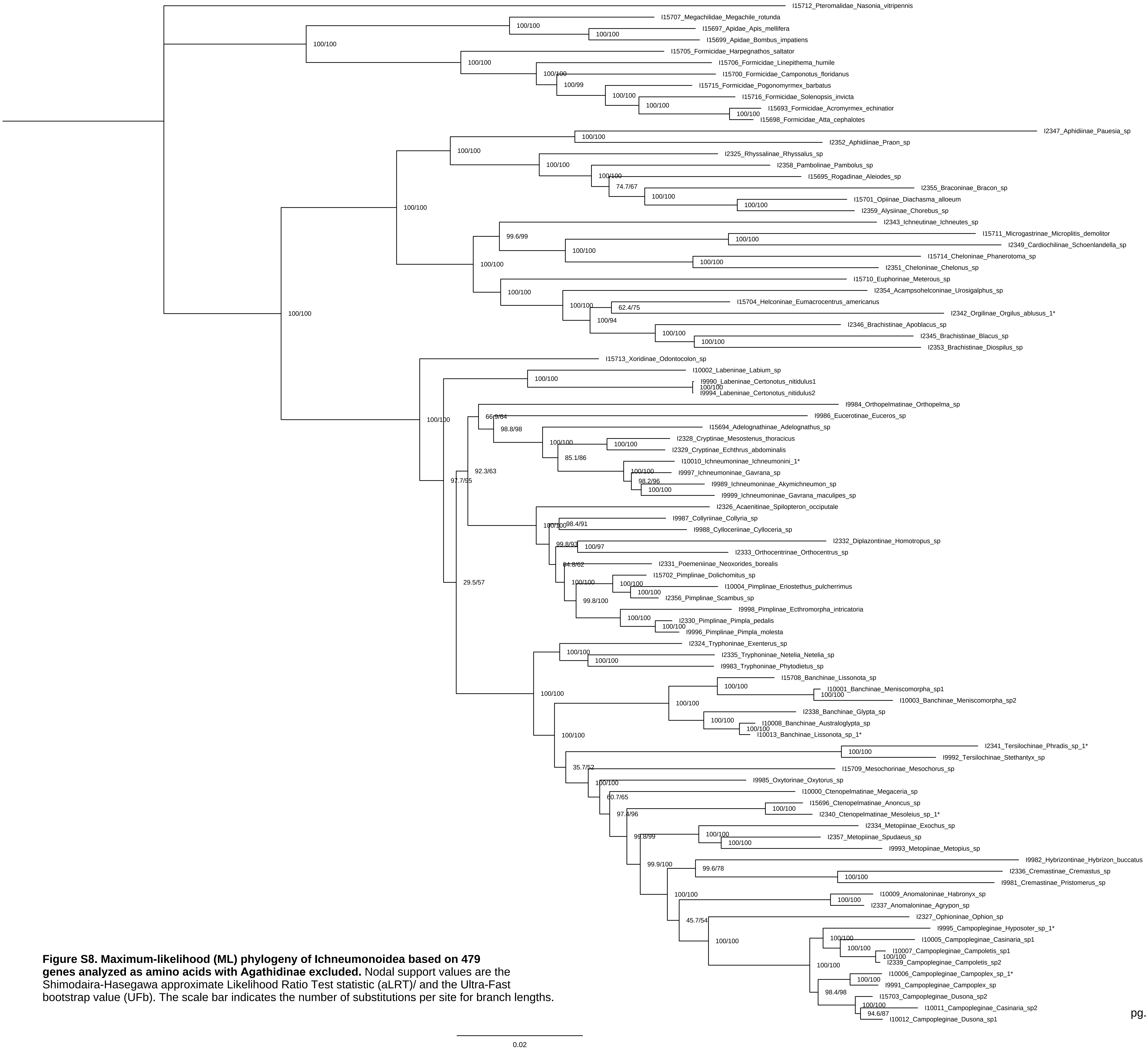


**Figure S8. Maximum-likelihood (ML) phylogeny of Ichneumonoidea based on 479 genes analyzed as amino acids with Agathidinae excluded.** Nodal support values are the Shimodaira-Hasegawa approximate Likelihood Ratio Test statistic (aLRT)/ and the Ultra-Fast bootstrap value (UFb). The scale bar indicates the number of substitutions per site for branch lengths.

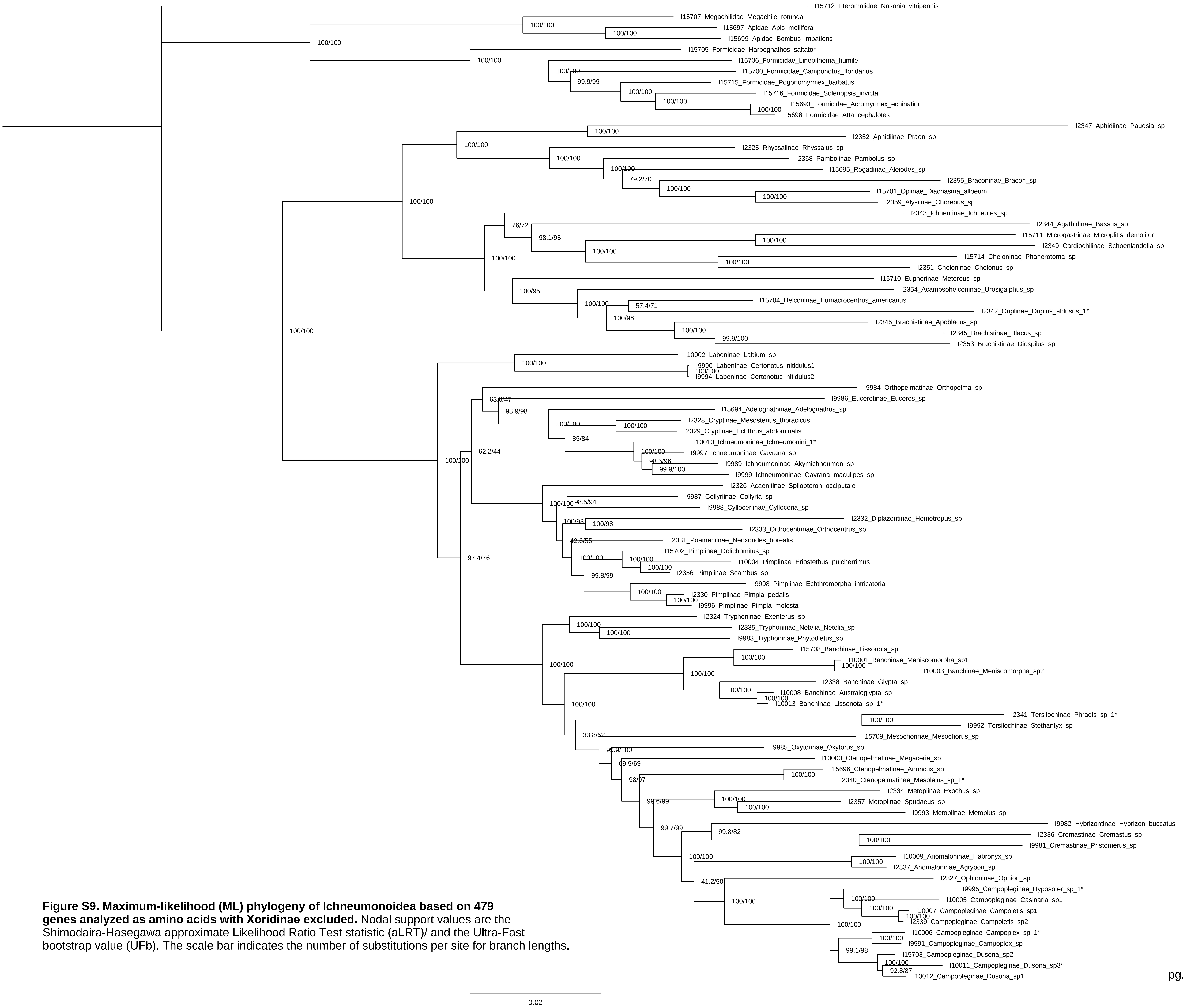


**Figure S9. Maximum-likelihood (ML) phylogeny of Ichneumonoidea based on 479 genes analyzed as amino acids with Xoridinae excluded.** Nodal support values are the Shimodaira-Hasegawa approximate Likelihood Ratio Test statistic (aLRT)/ and the Ultra-Fast bootstrap value (UFb). The scale bar indicates the number of substitutions per site for branch lengths.

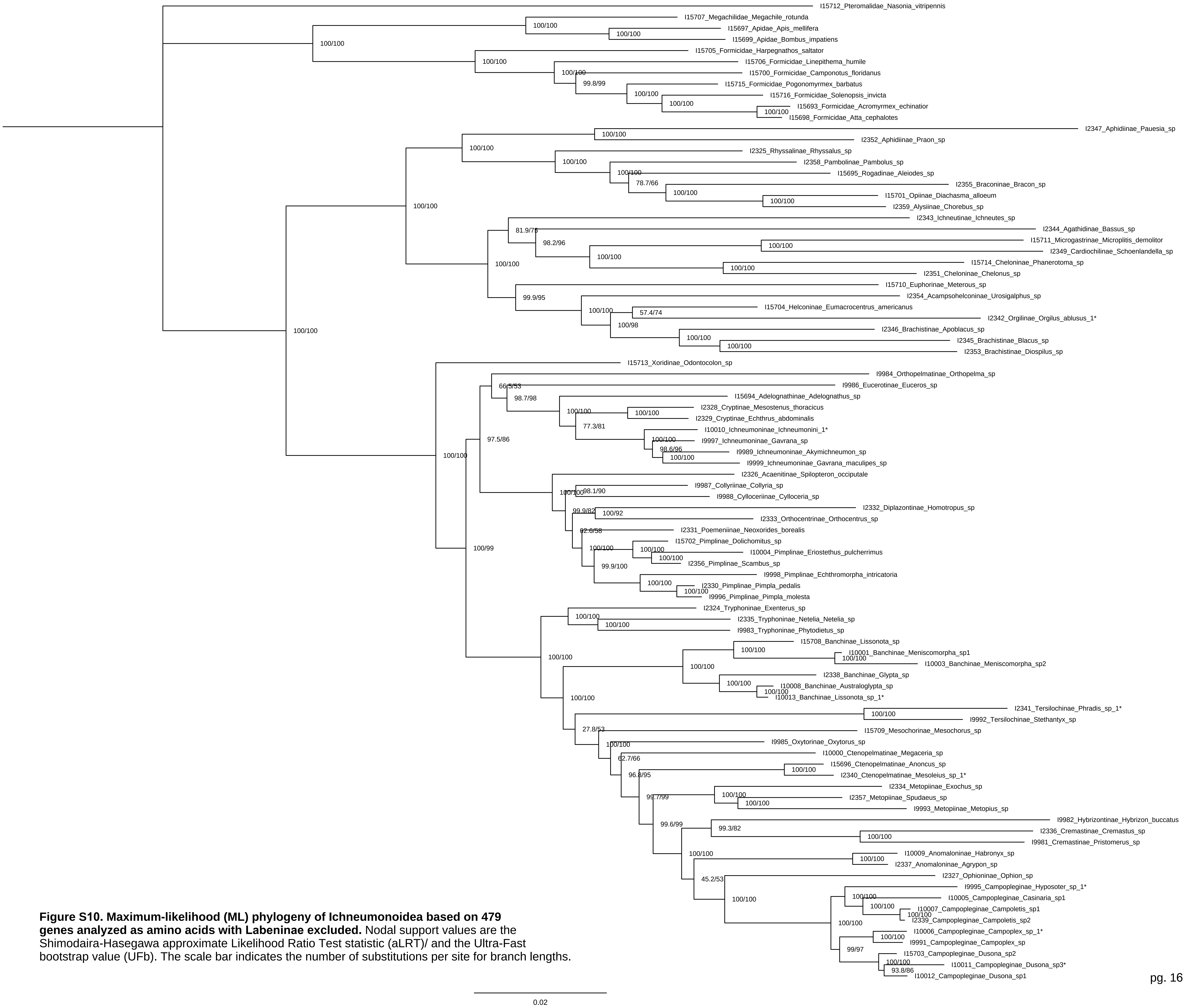


**Figure S10. Maximum-likelihood (ML) phylogeny of Ichneumonoidea based on 479 genes analyzed as amino acids with Labeninae excluded.** Nodal support values are the Shimodaira-Hasegawa approximate Likelihood Ratio Test statistic (aLRT)/ and the Ultra-Fast bootstrap value (UFb). The scale bar indicates the number of substitutions per site for branch lengths.

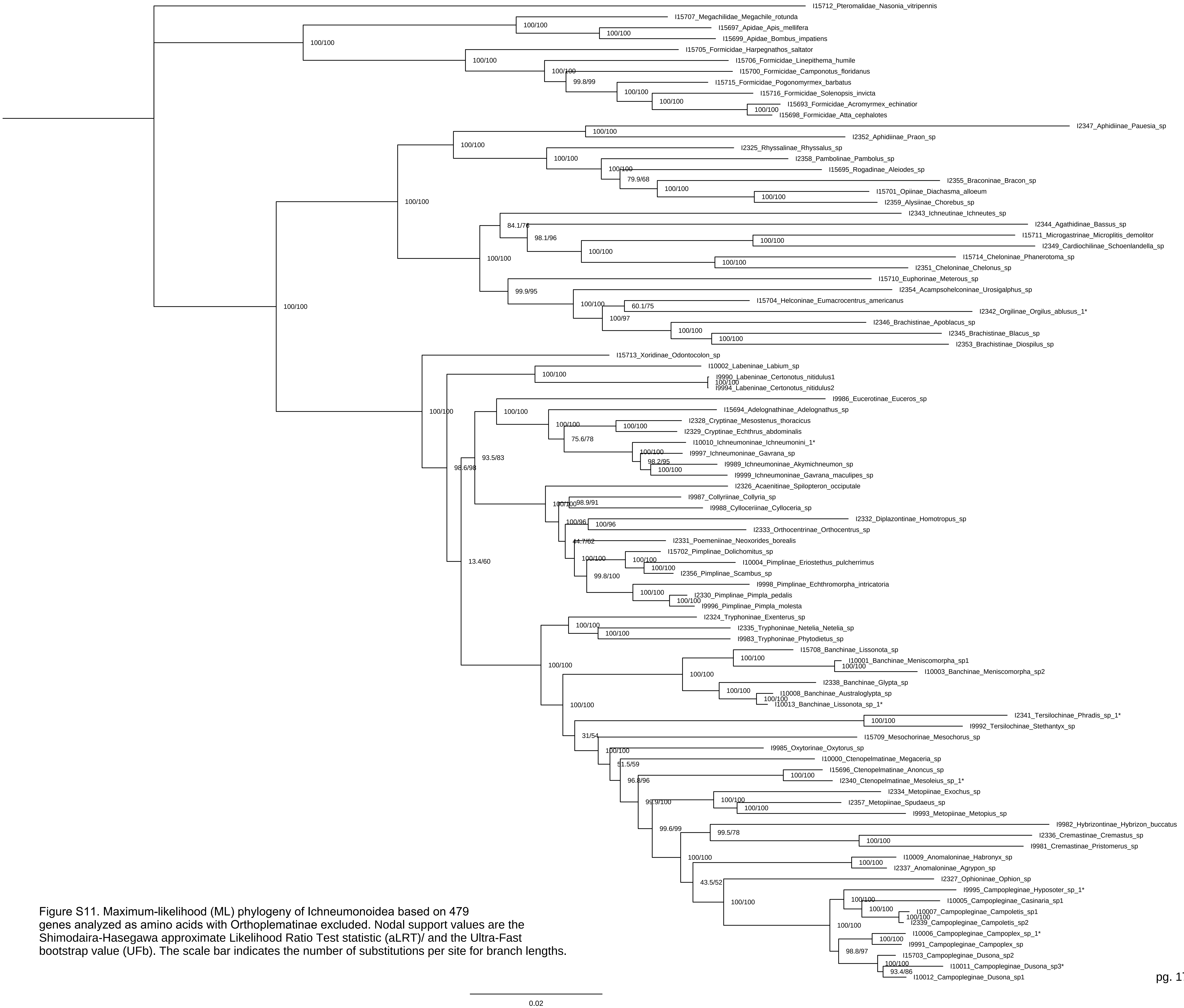


Figure S11. Maximum-likelihood (ML) phylogeny of Ichneumonoidea based on 479 genes analyzed as amino acids with Orthoplematinae excluded. Nodal support values are the Shimodaira-Hasegawa approximate Likelihood Ratio Test statistic (aLRT)/ and the Ultra-Fast bootstrap value (UFb). The scale bar indicates the number of substitutions per site for branch lengths.

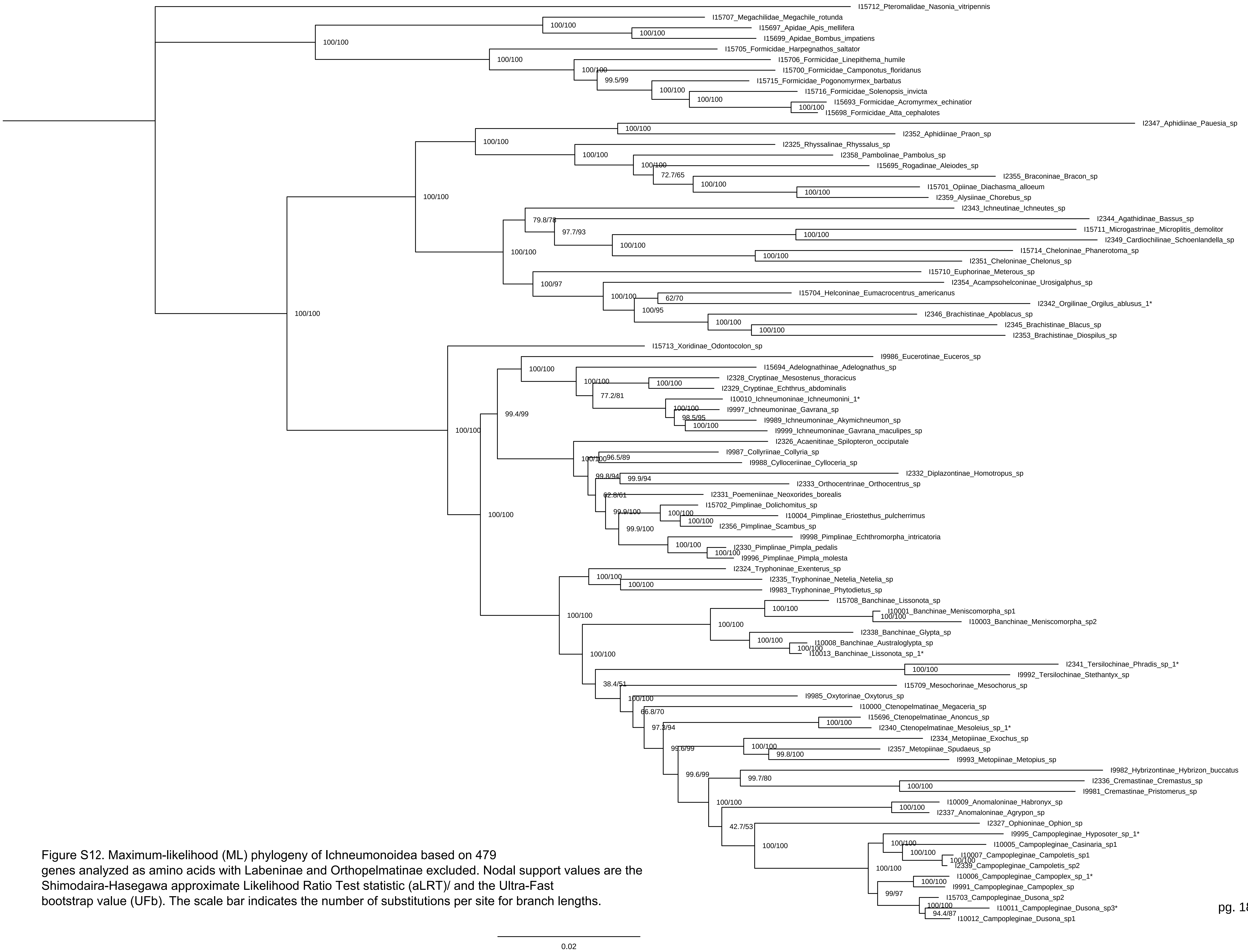


Figure S12. Maximum-likelihood (ML) phylogeny of Ichneumonoidea based on 479 genes analyzed as amino acids with Labeninae and Orthopelmatinae excluded. Nodal support values are the Shimodaira-Hasegawa approximate Likelihood Ratio Test statistic (aLRT)/ and the Ultra-Fast bootstrap value (UFb). The scale bar indicates the number of substitutions per site for branch lengths.

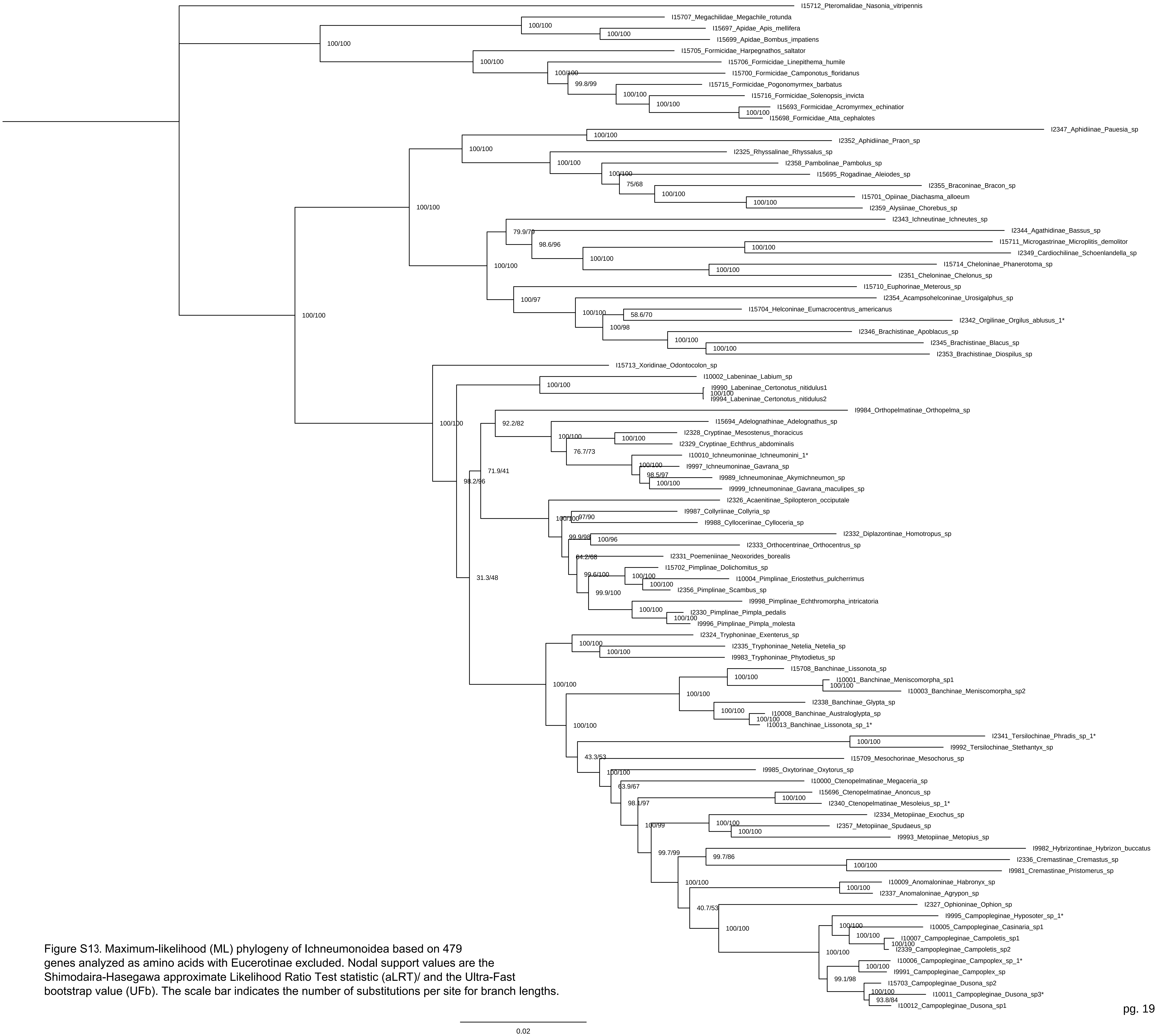


Figure S13. Maximum-likelihood (ML) phylogeny of Ichneumonoidea based on 479 genes analyzed as amino acids with Eucerotinae excluded. Nodal support values are the Shimodaira-Hasegawa approximate Likelihood Ratio Test statistic (aLRT)/ and the Ultra-Fast bootstrap value (UFb). The scale bar indicates the number of substitutions per site for branch lengths.

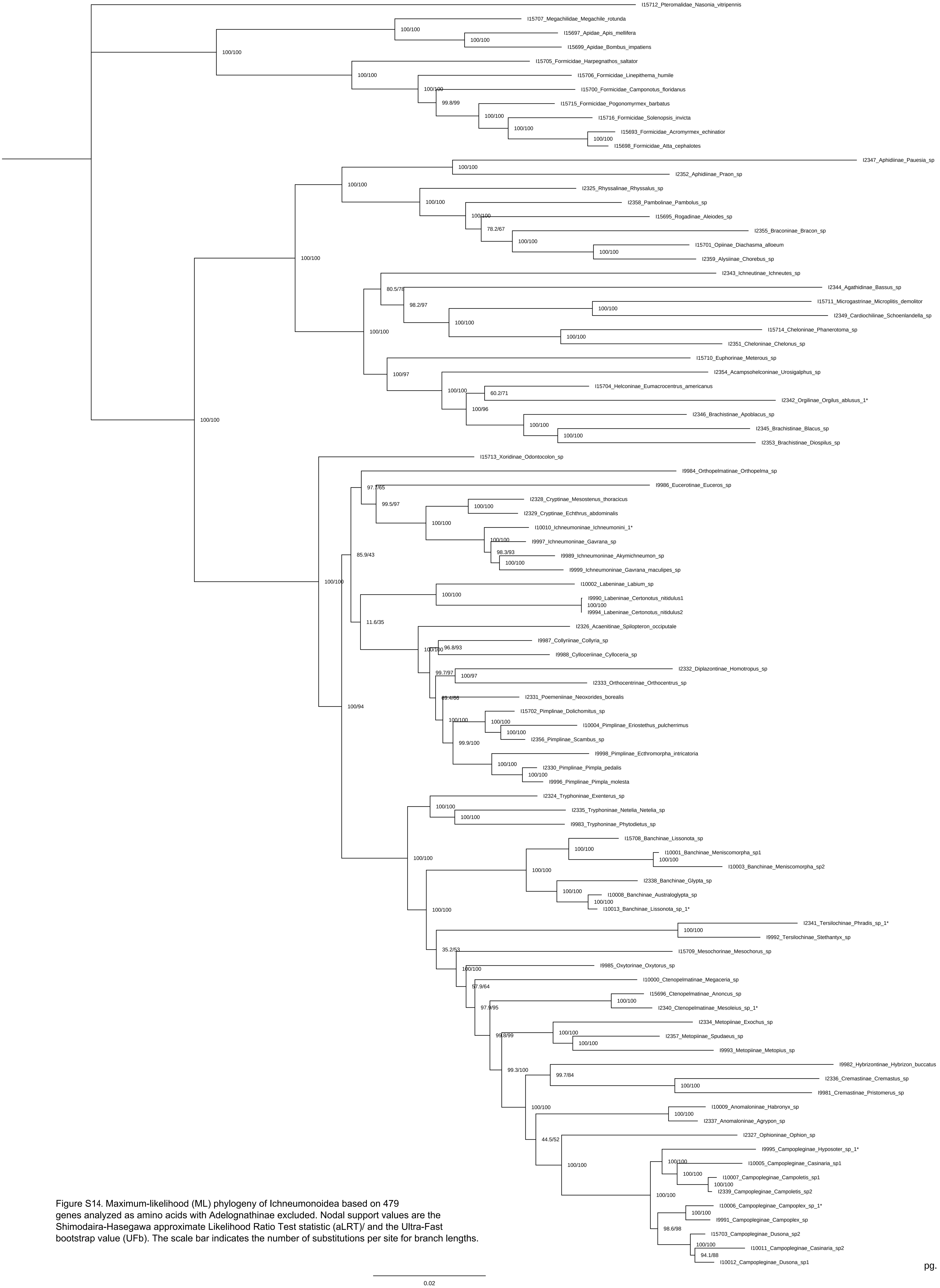


Figure S14. Maximum-likelihood (ML) phylogeny of Ichneumonoidea based on 479 genes analyzed as amino acids with Adelognathinae excluded. Nodal support values are the Shimodaira-Hasegawa approximate Likelihood Ratio Test statistic (aLRT)/ and the Ultra-Fast bootstrap value (UFb). The scale bar indicates the number of substitutions per site for branch lengths.

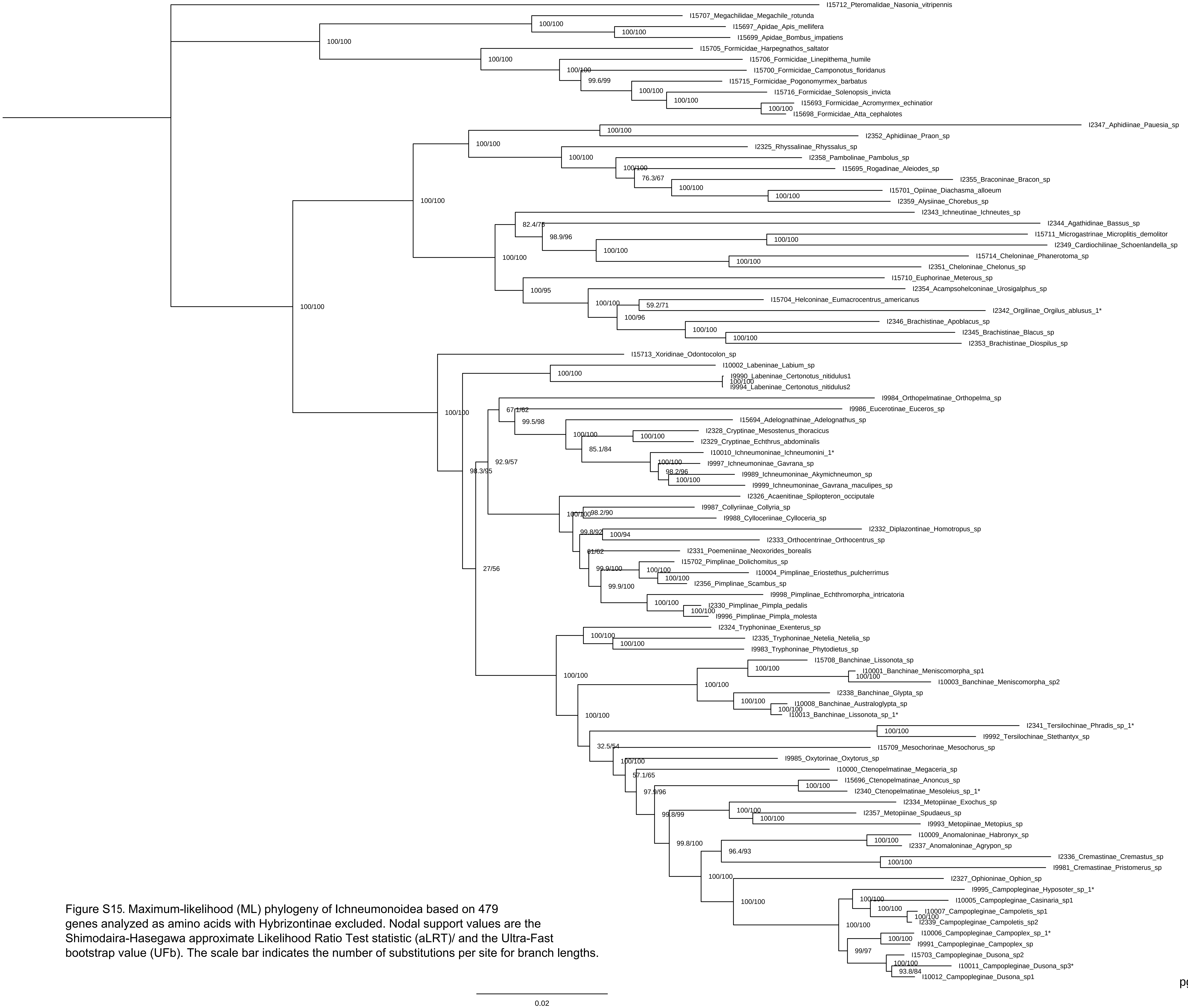


Figure S15. Maximum-likelihood (ML) phylogeny of Ichneumonoidea based on 479 genes analyzed as amino acids with Hybrizontinae excluded. Nodal support values are the Shimodaira-Hasegawa approximate Likelihood Ratio Test statistic (aLRT)/ and the Ultra-Fast bootstrap value (UFb). The scale bar indicates the number of substitutions per site for branch lengths.

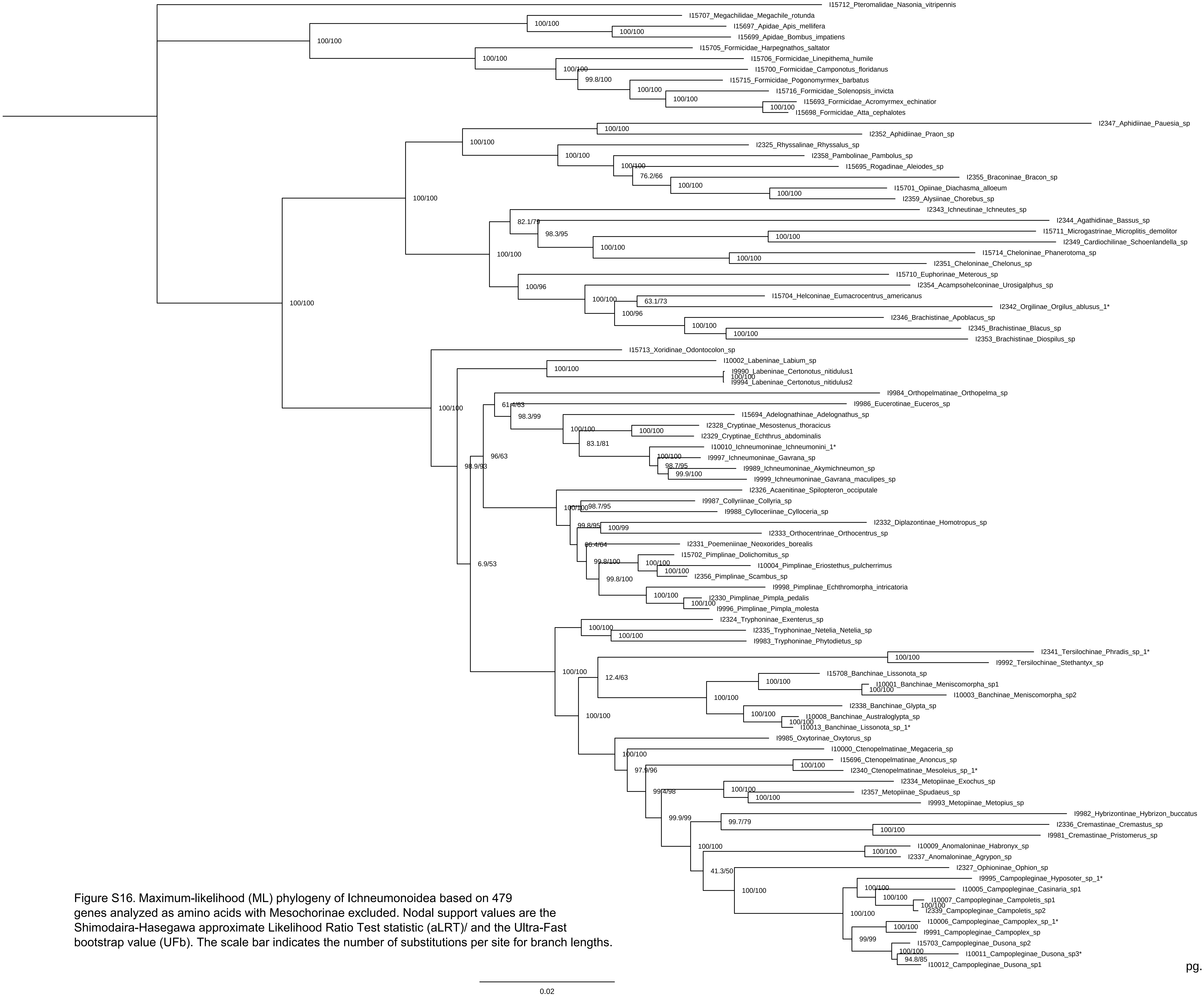


Figure S16. Maximum-likelihood (ML) phylogeny of Ichneumonoidea based on 479 genes analyzed as amino acids with Mesochorinae excluded. Nodal support values are the Shimodaira-Hasegawa approximate Likelihood Ratio Test statistic (aLRT)/ and the Ultra-Fast bootstrap value (UFb). The scale bar indicates the number of substitutions per site for branch lengths.

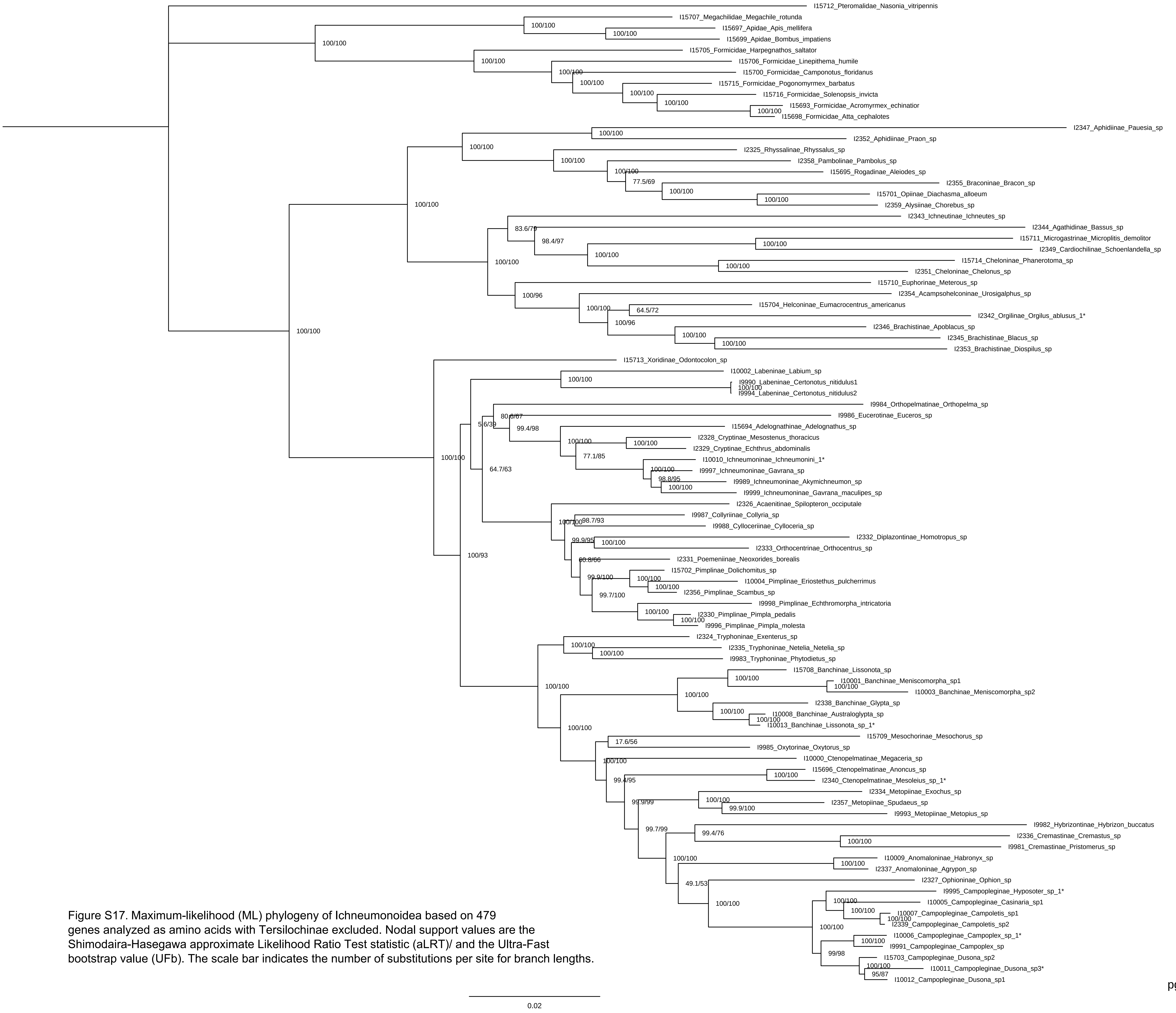


Figure S17. Maximum-likelihood (ML) phylogeny of Ichneumonoidea based on 479 genes analyzed as amino acids with Tersilochinae excluded. Nodal support values are the Shimodaira-Hasegawa approximate Likelihood Ratio Test statistic (aLRT)/ and the Ultra-Fast bootstrap value (UFb). The scale bar indicates the number of substitutions per site for branch lengths.

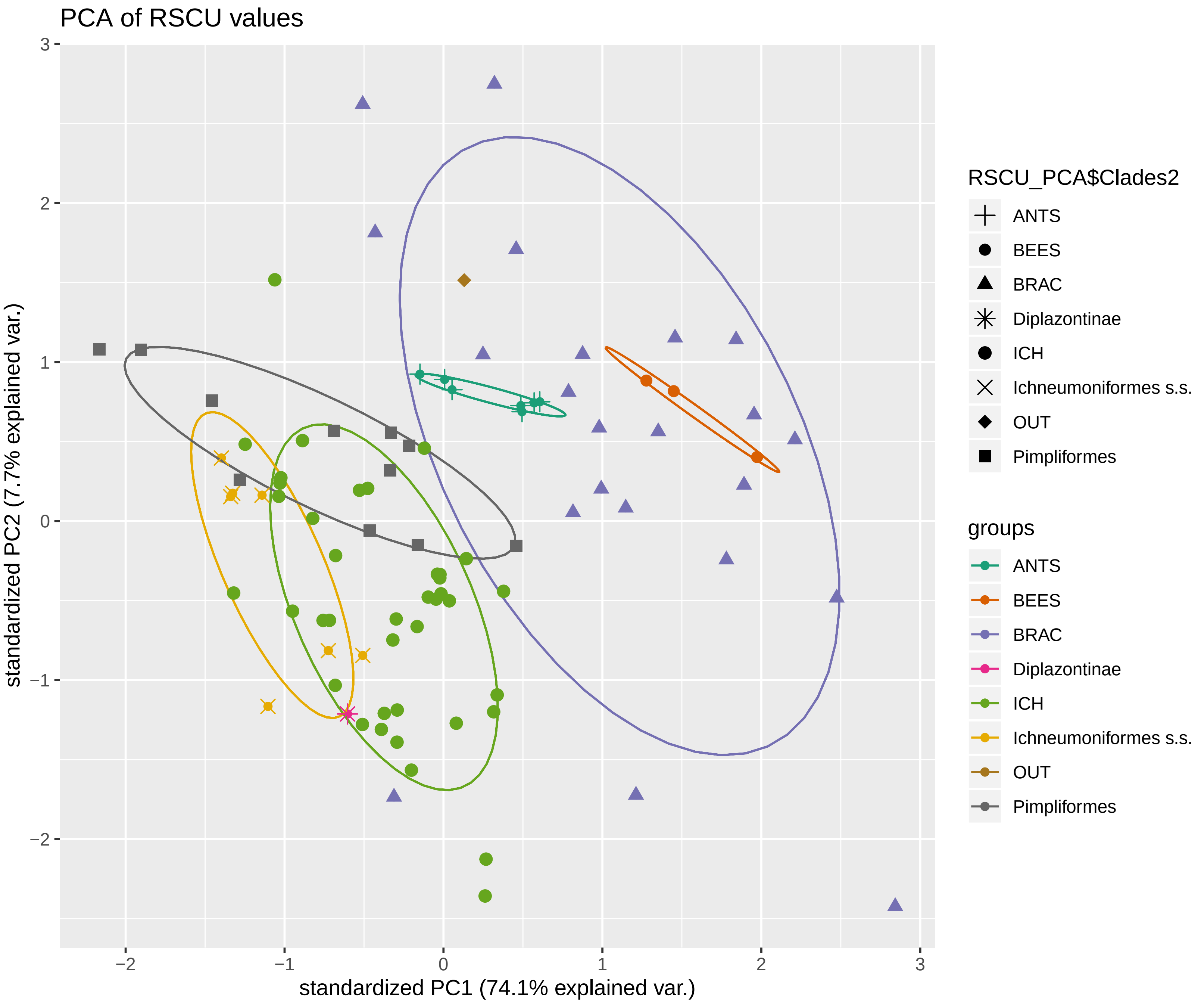


Figure S18. Principle Component Analysis (PCA) of Relative Synonymous Codon Usage (RSCU) between select major groups. OUT = Outgroups, Ants and Bees are outgroups; BRAC = Braconidae; ICH = Ichneumonidae. Diplazontinae is separated from Pimpliformes to show variation.

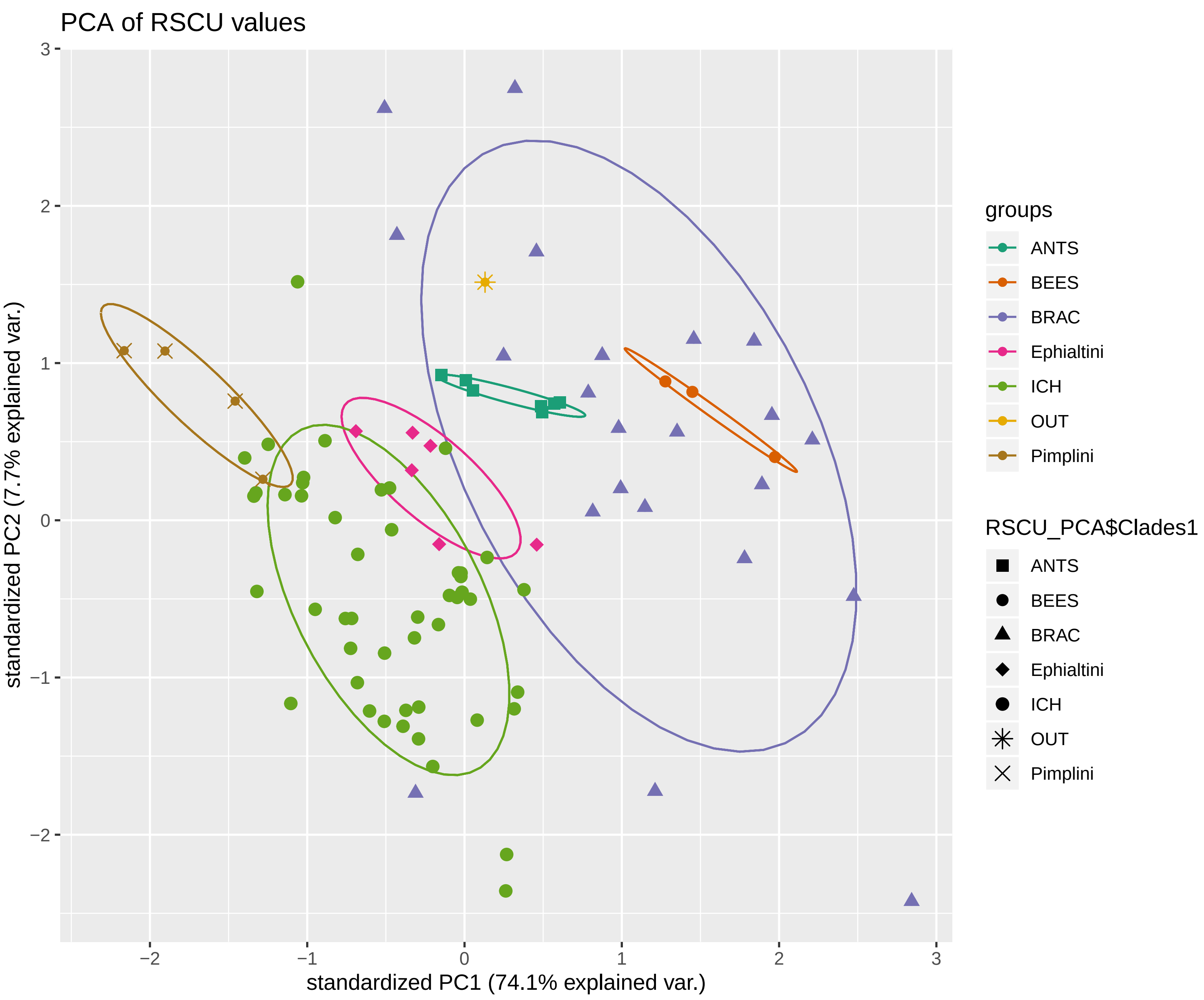


Figure S19. Principle Component Analysis (PCA) of Relative Synonymous Codon Usage (RSCU) between select major groups and sampled Pimplinae tribes (Ephaltini and Pimplini). OUT = Outgroups, Ants and Bees are outgroups; BRAC = Braconidae; ICH = Ichneumonidae. Diplazontinae is separated from Pimpliformes to show variation.
